# Pathophysiological remodeling of the skeletal muscle microenvironment in patients with lung cancer

**DOI:** 10.1101/2025.11.19.689342

**Authors:** Edmund Battey, Andrea Irazoki, Jakob Wang, Jonas Sørensen, Johanne L. Modvig, Scott Themsen, Christian T. Voldstedlund, Cathrine Knøs, Nicoline R. Andersen, Emma Frank, Søren Kudahl, Nanna Hahn, Anna Madsen, Steffen H. Raun, Clara Prats, Steen Larsen, Oksana Dmytriyeva, Niels Ørtenblad, Jean Farup, Lykke Sylow

**Author notes:** These authors contributed equally to this work (Edmund Battey, Andrea Irazoki, Jakob Wang). These authors jointly supervised this work (Jean Farup, Lykke Sylow).

## Abstract

**Background:** Muscle wasting, systemic inflammation, and functional decline are highly prevalent and detrimental in patients with advanced-stage non–small cell lung cancer (NSCLC).

**Methods:** In this cross-sectional study, we investigated NSCLC-associated muscle remodeling by analyzing skeletal muscle biopsies from patients with NSCLC (n = 18) and matched controls (n = 18) using quantitative proteomics, histology, fluorescence-activated cell sorting, gene expression profiling, and high-resolution respirometry.

**Findings:** NSCLC muscle was characterized by type II muscle fiber atrophy, greater collagen deposition, and redistribution of lipids to the extracellular matrix (ECM), together with remodeling of the inflammatory, immune, ECM and mitochondrial proteome. Additionally, mitochondrial respiratory capacity and morphology were altered in patients with NSCLC, which was associated with increased oxidative stress and dysregulated calcium handling. Concomitantly, we detected STAT3 activation and immune cell alterations, which may negatively impact skeletal muscle health in patients with NSCLC. Finally, we identified a shift in fibro-adipogenic progenitors (FAPs), favoring the CD90^−^ subtype. Mechanistically, conditioned media from patient-derived FAPs reduced myotube width in vitro, uncovering a novel mechanism by which altered paracrine signaling from the muscle-resident stromal compartment drives atrophy in cancer cachexia.

**Interpretation:** These findings provide human evidence that altered FAP composition, mitochondrial homeostasis, calcium handling, and immune cell landscape accompany muscle wasting in NSCLC, which may inform therapeutic strategies to preserve skeletal muscle health in patients with cancer.

**Funding:** Danish Diabetes and Endocrine Academy, Novo Nordisk Foundation, Independent Research Fund Denmark, Danish Cancer Society, Sejer Persson and Lis Klüwer Perssons Legat, A.P. Møller Foundation for the Advancement of Medical Science, Danish Medical Society Research Foundation, and Lundbeck Foundation.

**Research in Context:** *Evidence before this study:* We searched PubMed from database inception to September 1, 2026, with no language restrictions, using combinations of the terms “non-small cell lung cancer”, “lung cancer”, “cancer cachexia”, “cancer-induced muscle wasting”, “skeletal muscle atrophy”, “muscle fiber”, “fibrosis”, “fibro-adipogenic progenitors”, “mitochondria”, “calcium handling”, “immune cell”, “inflammation”, and “proteomics”, and we hand-searched the reference lists of relevant articles and reviews. While comprehensive human evidence remains exceedingly scarce, prior research has established several key components of cancer-associated muscle pathology: studies in patients with non-small cell lung cancer have described muscle fiber atrophy, systemic and local inflammation, altered muscle metabolic phenotypes, and changes occurring during anticancer treatment. Studies in other cancer populations and in experimental models of cancer cachexia have additionally implicated fibrosis, fatty infiltration, immune remodeling, and fibro-adipogenic progenitors. However, the available human studies were generally small and focused on individual components of muscle pathology, and we found no study integrating structural, proteomic, immune-cell, mitochondrial, calcium-handling, and fibro-adipogenic progenitor phenotypes in skeletal muscle from newly diagnosed patients with non-small cell lung cancer, or directly testing whether secreted factors from patient-derived fibro-adipogenic progenitors influence myotube size.

**Added value of this study:** Using skeletal muscle biopsies from newly diagnosed patients with advanced-stage non-small cell lung cancer and matched controls, this study provides an integrated characterization of the structural, molecular, and cellular muscle microenvironment. We identify preferential type II fiber atrophy, fibrosis, and extracellular lipid redistribution alongside inflammatory and immune remodeling, altered mitochondrial respiration and morphology, oxidative stress, and impaired sarcoplasmic-reticulum calcium handling, together with a shift toward CD90^-^ fibro-adipogenic progenitors. Conditioned media from patient-derived fibro-adipogenic progenitors reduced myotube diameter, providing evidence that altered stromal-cell secretory function may contribute directly to muscle fiber atrophy in human cancer.

**Implications of all the available evidence:** These findings extend the biological understanding of cancer-associated muscle deterioration beyond muscle fiber catabolism alone and identify coordinated remodeling of the wider skeletal muscle microenvironment in non-small cell lung cancer. The altered phenotype and secretory activity of fibro-adipogenic progenitors, in particular, link stromal remodeling to muscle fiber atrophy. Defining the factors responsible for this altered intercellular signaling, and whether they can be therapeutically modified, could reveal strategies to preserve skeletal muscle mass and function in patients with cancer.

## Introduction

Cancer-associated cachexia is a multifactorial syndrome characterized by unintentional weight loss and skeletal muscle wasting, contributing to poor prognosis^1–4^. Cancer-induced muscle wasting affects 30-60% of patients with non-small cell lung cancer (NSCLC)^5^. For these patients, loss of muscle mass and function severely impairs survival, tolerance to anticancer therapy, physical function, and quality of life^6^. Defining the molecular and cellular features of NSCLC-related muscle deterioration may enable the development of targeted therapies to improve treatment and survival.

Although cancer-associated cachexia is clinically defined around progressive weight loss and skeletal muscle loss, cachexia status does not necessarily capture cancer-related skeletal muscle pathology. The Fearon consensus criteria classify patients using weight loss, BMI and low muscle mass, whereas direct muscle phenotypes such as muscle fiber size, tissue composition, mitochondrial biology, inflammatory signaling and proteomic remodeling may vary within and across these clinical strata^4–8^. This is particularly relevant in heterogeneous patient cohorts, where muscle abnormalities may precede or occur independently of overt cachexia classification. Direct comparison of skeletal muscle from patients with NSCLC and controls may therefore reveal tissue and molecular remodeling across the clinical spectrum of wasting.

Within the skeletal muscle niche, certain cellular and molecular alterations in muscle fibers and adjacent cell types are likely to drive atrophy and degeneration upon cancer and anti-cancer treatment, while others act to preserve muscle mass and resist degeneration. The presence of immune cells and a pro-inflammatory environment within the skeletal muscle niche is key for remodeling and regeneration^9^. Certain immune cell subsets positively correlate with muscle mass and muscle fiber size in patients with cancer^10,11^. On the other hand, through the chronic secretion of pro-inflammatory cytokines such as interleukin-6 (IL-6) and tumor necrosis factor-alpha (TNF-α), immune cells may promote catabolic pathways, induce apoptosis and inhibit muscle regeneration, and ultimately, accelerate cancer-induced muscle wasting^12–14^. Alterations in skeletal muscle bioenergetics are also known to contribute to muscle atrophy in preclinical models of cancer-induced muscle wasting ^15–17^ and to induce inflammatory processes in muscle cells that result in immune cell infiltration in the tissue^18^. However, significant and consistent clinical evidence supporting these observations is currently lacking.

Maintaining functional interactions between fibro-adipogenic progenitors (FAPs) and muscle stem cells (MuSCs) may be crucial for avoiding cancer-induced pathological remodeling in skeletal muscle^19,20^. FAPs are mesenchymal progenitor cells residing in the skeletal muscle with the capacity to differentiate into fibrogenic or adipogenic lineages depending on subcellular phenotype and the surrounding factors of the muscle microenvironment^21,22^. Fibrosis is a common feature in rodent models of cancer-induced muscle wasting and has been found in patients with pancreatic cancer^23,24^. FAPs can also drive deposition of ectopic adipocytes in the extracellular matrix (ECM), which may directly contribute to the pathophysiology of cancer-induced muscle wasting^21,25–27^. Furthermore, FAPs are emerging as crucial and targetable mediators of intercellular cross-talk within the muscle in health and disease^28,29^, with preclinical models emphasizing their contribution to muscle atrophy in sarcopenia, denervation^30,31^, and inflammation^32^. However, to our knowledge, no human studies have directly examined the role of FAPs in skeletal muscle deterioration in cancer-induced muscle wasting.

Mass spectrometry (MS)–based proteomics have emerged as powerful tools for unbiased, system-wide quantifications, including broad proteome alterations of specific proteins or pathways^33^. Unravelling the proteomic changes in skeletal muscle from patients with cancer could provide important new insights into molecular changes and elucidate new targets to prevent pathological remodeling in cancer. Yet, how cancer influences the structural and cellular remodeling of the muscle microenvironment is unclear, and human molecular proteomics data are lacking^34,35^.

Here, using a combination of proteomics, fluorescence-activated cell sorting, histological and biochemical analyses, and *in vitro* experiments, we comprehensively characterized the skeletal muscle microenvironment in patients with advanced-stage NSCLC and matched controls. Patients with NSCLC displayed type II fiber atrophy, fibrosis and elevated ECM lipid deposition at the time of diagnosis, evident across a cohort spanning non-CAC, pre-CAC and CAC classifications. These alterations were accompanied by a shift towards an inflammatory phenotype, as well as shifts in immune cell composition, mitochondrial abnormalities, oxidative stress and calcium dysregulation. We also identified marked differences in FAP subpopulation composition uncovering a potential new mechanism by which paracrine signaling from the muscle-resident stromal compartment causes cachexia-related muscle atrophy . Our findings identify immune cell, mitochondrial and FAP remodeling as features of the skeletal muscle microenvironment in advanced-stage NSCLC, mechanistically highlighting potential therapeutic targets to mitigate muscle atrophy in these patients. Moreover, we provide an open-source NSCLC human muscle proteome resource, offering insights into the pathophysiology of NSCLC-induced muscle remodeling.

## Methods

### Participants

This cross-sectional observational study compared skeletal muscle from newly diagnosed patients with advanced-stage NSCLC with matched non-cancer controls. Participant characteristics are presented in Table 1, and the NSCLC cohort has been described in detail previously^36,37^. Both male and female participants were included in this study. Control participants were matched to patients with NSCLC by sex where possible, and sex distribution is reported in Table 1. Because of the limited sample size and the primary focus on NSCLC-associated skeletal muscle remodeling, the study was not powered to assess sex-specific effects. Accordingly, data from male and female participants were analyzed together, and potential sex-dependent differences should be investigated in larger future cohorts. The study size was determined by the availability of skeletal muscle biopsy material from 18 patients with NSCLC and 18 matched controls included in the present analysis; no separate a priori sample-size calculation was performed for these exploratory tissue analyses.

**Table 1.** Participant characteristics.

| Characteristic | Patients (N=18) | Controls (N=18) |
| --- | --- | --- |
| **Sex** |  |  |
| Male, n | 9 | 9 |
| Female, n | 9 | 9 |
| **Age (years)** |  |  |
| Mean (SD) | 64.2 (10.5) | 64.8 (7.5) |
| Range | 46-80 | 54-79 |
| **BMI (kg/m²)** |  |  |
| Mean (SD) | 24.9 (5.6) | 25.2 (5.8) |
| Range | 16-41 | 20-45 |
| BMI <20 kg/m², n (%) | 3 (16.7) | 0 (0) |
| **Cachexia status at diagnosis** |  |  |
| Cachectic, n (%) | 8 (44.4) | 0 (0) |
| Pre-cachectic, n (%) | 4 (22.2) | 0 (0) |
| Non-cachectic, n (%) | 6 (33.3) | 0 (0) |

Within the first week after being diagnosed, patients with advanced stage NSCLC were included at the Department of Oncology, Copenhagen University Hospital – Rigshospitalet, Copenhagen, between August 2022 and October 2023. Inclusion criteria were: (1) age > 18 years, (2) pathologically confirmed NSCLC (3) referred for first-line palliative systemic anticancer therapy (4) having a staging/baseline computed tomography (CT) scan within 4 weeks of initiation of treatment, or a baseline scan planned within the first week of treatment, (5) being in Eastern Cooperative Oncology Group Performance Score (ECOG PS) 0-2, (5) having signed the informed consent form to the study. Exclusion criteria were: (1) any other known malignancy requiring active treatment, (2) palliative radiotherapy as primary treatment, (3) ECOG PS >2, (4) physical disabilities excluding physical testing, (5) inability to understand Danish, (6) inability to understand scoring systems/patient-reported outcome measures, (7) taking anti-coagulant therapy that could not be discontinued for the muscle biopsy. If the oncologist found a patient capable of receiving anticancer treatment, no other comorbidities were reason for exclusion. Patients were classified, according to self-reported weight loss over 6 months prior to their diagnoses as: CAC (more than 5% loss of stable body weight over the past 6 months), or a BMI less than 20 kg/m² and ongoing weight loss (WL) of more than 2%, or sarcopenia and ongoing WL of more than 2% but have not entered the refractory stage; pre-CAC (1-5% WL), or non-CAC (increase or up to 1% WL. Sarcopenia was defined as L3 CT-derived skeletal muscle index (SMI) (cm^2^/m^2^) < 43/41 for normal or underweight men/women and <53/41 for overweight and obese men/women) ^4,7^. 18 non-cancer and weight-stable control subjects, matched for age, sex, BMI and co-morbidity, were also included.

#### Body composition

Body composition was obtained using a dual-energy X-ray absorptiometry (DEXA) scan at the MuscleLab at Copenhagen University Hospital – Rigshospitalet.

#### Blood sample and muscle tissue collection

The day before the first systemic treatment was initiated and 12 weeks into first-line treatment non-fasted blood samples were taken. Analyses were performed at the Dept. of Clinical Biochemistry Copenhagen University Hospital - Rigshospitalet. In the same first week of systemic treatment (median day 6), the patients were invited to the oncological outpatient clinic after an overnight fast. Patients were urged not to do exercise 48 hrs prior to their biopsy visit.

Percutaneous biopsy of the left *vastus lateralis* was obtained from the third most proximal aspect of the patella and the inguinal fold under local anaesthetic (lidocaine without adrenalin 20mg/ml). A 6-mm incision was made through the skin, fat, and down to fascia, and the biopsy needle (6 mm Bergström) was advanced past the fascia into the muscle. Suction was applied and the tissue was excised.

#### Muscle tissue processing

All biopsies were rinsed in ice cold saline within 30 s after procedure, and within 1-2 min a portion of fresh *Vastus lateralis* was quickly snap frozen in liquid nitrogen and stored at −70 °C. Biopsies were pulverized using a Bessman type tissue pulverizer (Cellcrusher kit) on dry ice.

To preserve and assess peroxiredoxin 2 and 3 (PRDX2 and 3) dimerization ratios and H2O2 levels, a fresh portion of the muscle biopsy was incubated for 10 min in ice-cold 100 mM N-Ethylmaleimide (NEM) diluted in PBS. NEM was subsequently aspirated, and the tissue snap frozen.

For RNA extraction, samples were homogenized in QIAzol lysis reagent (#79306 Qiagen) with three sterile ceramic beads and using the TissueLyser II bead mill (Qiagen, USA) for 3 min at 30 Hz.

For protein extraction, tissues were homogenized in ice cold homogenization buffer (10% glycerol, 1% NP-40, 20 mM sodium pyrophosphate, 150 mM NaCl, 50 mM HEPES (pH 7.5), 20 mM β-glycerophosphate, 10 mM NaF, 2mM phenylmethylsulfonyl fluoride (PMSF), 1 mM EDTA (pH 8.0), 2 mM Na3VO4, 10 μg/mL leupeptin, 10 μg/mL aproptinin, 3 mM benzamidine, 100 mM NEM) with a steel bead using the TissueLyser II bead mill (Qiagen, USA) for 1 min at 30 Hz.

For flow cytometry, 80-200 mg of the muscle biopsies were immediately submerged in ice-cold Hams F10+ including bicarbonate and glutamine (#N6908, Sigma-Aldrich, Denmark), 10% horse serum (#26050088, Gibco, Thermo Fisher, MA) and 1% pen/strep (#15140122, Gibco, Thermo Fisher, MA) (wash-buffer). Samples were maintained on ice until further processing (within 1 hour), based on a previously described protocol^38^. Single cell suspensions were obtained via enzymatic digestion using 700 U/mL collagenase II (#46D16552, Worthington, Lakewood, NJ) and 3.27 U/mL dispase I (#04 942 078 001, Roche Diagnostics, Basel, Switzerland). After the addition of enzymes, samples were placed in a 37 °C water bath for 60 minutes and triturated with a pipette every 20 minutes to aid digestion. Enzymes were blocked through addition of wash-buffer, and the cell suspension was triturated through a 18G sterile needle 5 times and filtered through 70-um cell strainer (#130-110-916, Miltenyi Biotech). Finally, the cell suspension was centrifuged at 500 g for 5 min, before the supernatant was removed and the cell pellet was resuspended in 500 uL of cryopreservation buffer (StemMACs, #130-109-558 Miltenyi Biotech) and stored at −150 °C until the day of sorting. Prior to digestion, the sample weight was recorded to enable normalization of absolute cell counts to tissue weight.

#### Mass spectrometry and proteomics

Approximately 5-7 mg of skeletal muscle biopsy tissue was transferred to Twintec plates for proteomic analysis using the Astral platform. Raw data were processed without imputation of missing values prior to differential expression analysis to minimize artificial variance. To account for multiple hypothesis testing, permutation-based testing with 15,000 iterations was applied. Proteins with an adjusted P < 0.05 and log₂ fold-change > 0.25 or <-0.25 were taken forward for Gene Ontology (GO) over-representation analysis. Enrichment analysis was performed using STRING v12, with all detected proteins in the dataset used as the statistical background. For visualization, enriched GO terms were selected from the top five upregulated and top ten downregulated terms ranked by STRING “signal” score and presented as bubble plots. The signal score is calculated as a weighted harmonic mean of the observed/expected ratio and −log (FDR), and terms were selected to represent the dominant non-redundant biological themes. Gene Set Enrichment Analysis (GSEA) was performed using the R package clusterProfiler using Homo sapiens as reference organism, with all quantified proteins used as background.

#### Histology

Muscle biopsies were embedded in optimal cutting temperature (OCT) compound and cryosectioned at 8 μm thickness with fibers oriented perpendicular to the blade to ensure accurate assessment of muscle fiber morphology and size^39^.

For fiber typing and membrane delineation, sections were blocked in 5% donkey serum and incubated with primary antibodies for 3 hours at room temperature (Supplementary Table 1). Corresponding secondary antibodies were used: DyLight 488–conjugated anti–mouse IgG1 (Jackson #115-545-205), DyLight 405–conjugated anti–mouse IgM (Jackson #715-475-020), and Alexa Fluor 568–conjugated anti–rabbit (Invitrogen #A11011). Nuclei were counterstained with Sytox™ Deep Red (1:2000). Imaging was performed on a Carl Zeiss AxioObserver microscope through a Plan-Apochromat 20x/0.8 objective.

To assess lipid accumulation, sections were fixed in methanol-free 4% formaldehyde (Thermo Fisher Scientific #28908) for 20 minutes, followed by incubation with 0.5 µM BODIPY™ 493/503 (Invitrogen #D3922) and Sytox™ Deep Red nuclear stain (1:2000; Invitrogen #S11380) for 30 minutes at room temperature. Images were acquired using a Carl Zeiss AxioObserver microscope through a Plan-Apochromat 20x/0.8 objective.

Collagen content was evaluated using the Picrosirius Red staining kit (Abcam #ab150681). Sections were fixed in 4% formaldehyde for 20 minutes, washed in distilled water, and stained with Sirius Red for 60 minutes. Excess dye was removed by washing in 0.5% acetic acid, followed by dehydration, xylene clearing, and mounting in Pertex. Brightfield and polarized light images were acquired using the Carl Zeiss AxioScan 7, through a Plan-Apochromat 10x/0.45 objective.

Glycogen deposition was assessed performing periodic acid-Schiff (PAS) staining. Fresh frozen muscle sections were fixed in 4% formaldehyde for 20 minutes and then processed for PAS staining according to the manufacturer’s protocol (395B-KT, Sigma-Aldrich, Søborg, Denmark). The sections were treated with periodic acid solution for 5 minutes. After sequential washes in running tap water and distilled water, they were stained with Schiff reagent for 15 minutes, followed by an additional wash in running tap water. Finally, the sections were dehydrated, cleared with xylene, and mounted using Pertex. Brightfield images were acquired using a Zeiss AxioObserver microscope by a blinded investigator.

Quantification of collagen and lipid deposition was done with QuPath software^40^. The fractional area of polarized signal (Picrosirius Red) was measured by thresholding and normalizing it to the whole-tissue section. Lipid content was analyzed by thresholding the BODIPY-positive pixels in five fields per section, and fiber-specific signal intensity was quantified from 50–100 randomly chosen individual fibers using Fiji^41^. Membrane-to-cytosol lipid stain intensity ratios were calculated by thresholding (MinError) laminin-defined membranes and generating region of interests, which were transferred to the lipid images for intensity measurements. Muscle fiber cross-sectional area and fiber type distribution were determined using the Cellpose extension in QuPath on whole-section images based on MyHC staining profiles.

#### Myosin Heavy Chain composition analysis

Myosin heavy chain isoform composition was analyzed in muscle biopsy samples using large-format SDS-PAGE to achieve high-resolution separation of isoforms. Muscle homogenates were prepared and loaded in duplicate lanes at two different protein concentrations to ensure linear detection. Samples were electrophoresed on 6% polyacrylamide gels under denaturing conditions for 68 h at low voltage, allowing clear resolution of I, IIA and IIX isoforms. Gels were subsequently coomassie-stained, and band intensities were quantified densitometrically. The relative abundance of each MHC isoform was expressed as a percentage of total myosin heavy chain content.

#### Respirometry

A portion of fresh muscle was placed in ice-cold BIOPS solution (OROBOROS Instruments, Innsbruck, Austria) immediately after dissection. Muscle fibers were mechanically separated with needles on ice. Fibers were then permeabilized with 50 μg/mL saponin for 30 min on ice with shaking, followed by two 10 min washes in MIRO5 respiration medium (OROBOROS Instruments, Innsbruck, Austria). Approximately 2-3 mg of muscle fibers were transferred to the oxygraph-2k (OROBOROS Instruments, Innsbruck, Austria) containing MIRO5 respiration medium. Respiration measurements were performed at 37 °C in hyperoxia ([O2: 200-450 μmol/L). The protocol included adding 10 mM glutamate, 2 mM malate, and 5 mM sodium pyruvate to assess resting respiration (state II, leak respiration), followed by 2.5 mM ADP for state III. Succinate at 10 mM was added to measure full electron transport chain (ETC) capacity, upon which complexes I and III were inhibited by adding 0.5 μM rotenone and 2.5 μM antimycin A.

#### Single fiber isolation and immunostaining

Skeletal muscle biopsy samples were fixed in Zamboni buffer (2% paraformaldehyde supplemented with 1.2% picric acid) for 30 min at room temperature, followed by 4 hours in fresh fixative at 4 °C^42^. Biopsies were then dissected into bundles of 10–20 fibers, transferred to 1:1 PBS/glycerol (v/v) for 1-3 days, and stored at −20 °C. Fibers were further dissected into bundles of approximately 2-5 fibers. After permeabilizing the muscle fibers with 0.2% Triton x100 and blocking non-specific binding sites with 5% goat serum and, muscle fibers were stained to visualize type I fibers and mitochondria using primary antibodies that detect Myosin Heavy Chain 7 (Developmental Studies Hybridoma Bank, AB_10572253) and COXIV (Abcam, ab16056), respectively. The organization of intermyofibrillar mitochondria was determined by thresholding the COXIV signal in ImageJ^41^. Mitochondrial fractional area (%) was determined by calculating the area of the region of interest occupied by the COXIV signal divided by the total area of the region of interest. Fragmentation index was calculated by dividing the number of mitochondrial objects by the area of the mitochondria signal. 17.7 ± 3.8 fibers were analyzed per biopsy by a blinded investigator.

### Gene expression analyses

#### Gene expression in skeletal muscle biopsies

RNA was extracted using the RNeasy Mini Kit (Qiagen #74106) per the manufacturer’s instructions. RNA purity was assessed with the NanodropTM 2000/20000c spectrometer and ND1000 software (ThermoFisher Scientific). Reverse transcription was performed with 4 μL of qScript Ultra SuperMix (#95217, Quantabio) per 500 ng of RNA. Gene expression was quantified by real-time quantitative PCR using SYBR green (Applied Biosystems) and the QuantStudio 6 and 7 Real-Time PCR System (Applied Biosystems), following the default comparative Ct protocol. Measurements were normalized to a housekeeping gene (*RPLP0*). Primers and their sources are listed in Supplementary Table 2.

#### Single-cell RNAseq (10x Genomics)

Single cell gene expression data was obtained from our publicly available dataset deposited at the European Genome Archive (EGA number: EGAD00001008126). Detailed description of the sequencing procedures are previously reported elsewhere^43^.

#### Muscle H2O2 quantification

The Amplex™ Red Hydrogen Peroxide/Peroxidase Assay Kit (Thermo Fisher) was used following the manufacturer’s indications.

#### Immunoblotting

After tissue homogenization for protein extraction, samples were rotated end-over-end for 30 min at 4 °C, then centrifuged at 13000 x g for 15 min at 4 °C. Supernatants were collected and stored at −80 °C. Protein concentration was measured using the BCA method (Thermo Fisher #23225) with BSA as standard (Sigma-Aldrich #P0834). Protein extracts were resolved in 7%, 10%, or 12.5% Mini-PROTEAN precast acrylamide gels (BioRad) for SDS-PAGE, transferred to PVDF membranes, and blocked in TBS-Tween 20 with 2% milk protein for 5 min at room temperature. Membranes were incubated with primary antibodies overnight at 4 °C, followed by HRP-conjugated secondary antibodies for 45 min at room temperature. Pierce™ Reversible Stain (Thermo Fisher #24585) was used as loading control. Bands were visualized using the Bio-Rad ChemiDoc MP Imaging System and ECL+ (Amersham Biosciences). Band densitometry was performed with Quantity One (Bio-Rad) and bands were related to their own protein staining to correct for uneven loading. Primary antibodies are listed in Supplementary Table 1. When analyzing dimerization ratios of PRDX2 and PRDX3, protein extracts were prepared in the absence of the reducing agent DTT.

#### Calcium uptake and release assays

Calcium uptake and release rates in sarcoplasmic reticulum (SR) vesicles were measured using a fluorescent dye technique. Whole tissue homogenates were prepared from frozen specimens using a Potter–Elvehjem glass homogenizer (Kontes Glass Industry, Vineland, NJ) in an ice-cold buffer at a wet weight-to-volume ratio of 1:10. The buffer composition included 300 mM sucrose, 1 mM EDTA, 10 mM NaN₃, 40 mM Tris base, and 40 mM L-histidine, adjusted to pH 7.8, as described in detail elsewhere^44,45^. Calcium levels were measured using the fluorescent indicator Indo-1 (1 μM) at a sampling frequency of 20 Hz with a Ratiomaster RCM system (Photon Technology International, Brunswick, NJ, USA). Calcium uptake supported by oxalate was initiated by the addition of 2 mM ATP to reach a final concentration of 5 mM. Uptake was recorded for approximately 3 minutes until calcium levels plateaued below 100 nM, with no significant differences in plateau observed between groups.

The SR calcium uptake rate, referred to as Tau, was defined as the time constant of the single-exponential fit (the time required for free calcium concentration to decline by 63%, equivalent to 1/e of the initial value)^46^. Uptake rates were also determined at two calcium concentrations: 400 nM (high) and 200 nM (low), corresponding to muscle activation and resting levels of calcium. Following uptake measurements, cyclopiazonic acid was used to inhibit the SR calcium ATPase to estimate vesicular calcium leak. SR calcium release was then triggered by the addition of 4-Chloro-m-cresol (5 mM).

Calcium data were mathematically analyzed using monoexponential curve fitting (Curve Fitting Toolbox version 1.1.1; The MathWorks, Natick, MA, USA), as previously described^45^. Calcium uptake and release rates are expressed in relative terms as nmol calcium per gram wet weight per minute (nmol·g^−^¹ ww·min^−^¹). Tau is expressed reciprocally (s^−^¹·g^−^¹ ww), with higher values indicating faster calcium reuptake. All SR assays were conducted in duplicate, performed by a blinded investigator, and the results were averaged.

#### Plasma GDF-15 and CRP measurements

Fasting blood samples were drawn at 10 am on biopsy days. A 9 ml dry tube stood for a minimum of 30 min and maximum 1 hour at room temperature. Samples were then spun down at 2000×g for 10 min at 4°C. 500 μl serum was aliquoted and stored at −80°C. Plasma GDF-15 concentration was measured using a Human GDF-15 Quantikine ELISA kit (R&D Systems, #DGD150).

Plasma C-reactive protein (CRP) concentrations were analyzed using a latex particle–based immunoassay (turbidimetry) on a Cobas 8000 analyzer (c702 module, Roche Diagnostics). The method is standardized against the certified reference material ERM-DA474/IFCC, with a measurement range of 1–700 mg/L and a reference interval of <10 mg/L.

#### Fluorescence-activated cell sorting (FACS)

On days of cell sorting, frozen vials with the single cell suspensions were briefly thawed in a 37°C water bath and transferred to tubes containing 10 mL of wash buffer. Each vial was washed three times to optimize cell yield, before the tube was centrifuged at 500 g for 5 min at room temperature. Following aspiration of the supernatant, samples were resuspended in 150 μL wash buffer containing a mixture of antibodies (Supplementary Table 3).

Samples were protected from light to avoid bleaching and incubated during mild agitation for 30 minutes at 5°C. Following incubation, the antibody mixture was diluted with 15 mL wash buffer and centrifuged at 500 g for 5 minutes at RT. The supernatant was removed, and the pellet was resuspended in 900 μL wash buffer prior to being transferred into 5 mL FACS tubes through 30 μm cell strainers. Cells remained on ice and were protected from light until FACS analysis.

Immediately before sorting propidium iodide and counting beads were added to each sample, for assessment of cell viability and absolute cell counts, respectively. Samples were briefly vortexed before sorting to loosen any precipitate. Sorting was performed on the Bigfoot cell sorter (Thermo Fisher Scientific) using the 100 μm nozzle at 20 PSI to minimize cellular strain. Flow data were collected in the Sasquatch software and subsequently analyzed using FlowJo (version 10.6.2), with the gating strategy being ascertained using fluorescence minus one controls. Freshly isolated FAPs and MuSCs were collected in wash buffer and plated for in vitro experiments as outlined below.

FAP and MuSC analyses were performed in a subgroup of seven participants per group for whom single-cell preparations were available for FACS and downstream functional analyses.

### In vitro studies

For assessment of proliferation capacity of FAPs and MuSCs, we utilized the EdU incorporation assay as previously described^43^. Freshly isolated FAPs and MuSCs were seeded in 96-well half area plates (Corning, NY) pre-coated with ECM (ECM gel, #E1270, Sigma). Cells were seeded at an appx. density of 10000 and 6000 cells/well in case of FAPs and MuSCs, respectively. Cells were seeded in 180 uL of wash-buffer including 10 uM of the EdU nucleotide. Hereafter, cells were left in the incubator untouched until the time fixation; three and five days following FACS in case of MuSCs and FAPs, respectively. Prior to fixation (4% PFA for 5 minutes) the media was collected and stored at −80 °C until further used in conditioned-media experiments as described below. The wells were scanned using the EVOS M7000 imaging system (Thermo Fisher Scientific), and stitched images covering the entire well were then used to quantify the proportion of proliferating cells using ImageJ (NIH).

To investigate the potential effect of secreted factors from FAPs on myotube size regulation, we utilized conditioned media from the EdU-experiments in a C2C12-derived myotube cell culture. Here, C2C12 myoblasts (#91031101, Sigma Aldrich, passage 18) were seeded at 3000 cells/well in 96 well plates following cell expansion in a T75 culture flask. Cells were then grown to 70-80% confluency in high glucose (4.5 g/L) DMEM including 20% fetal bovine serum and 1% pen/strep, prior to initiation of differentiation by serum starvation using 2% horse serum in low-glucose DMEM. Media was changed every other day for 12 days. During the 48 hours of FAP media exposure, the differentiation media was supplemented with 50% conditioned media from the EdU culture, with a replenishment of the media following the first 24 hours of incubation. Following fixation, myotubes were stained for myosin heavy chain (MF20) and counterstained with DAPI. Images were acquired of the entire well using the EVOS M700 automated imaging system (Thermo Fisher Scientific). Myotube size was assessed as the average diameter of three randomly selected measurements along the length of each myotube, and the mean myotube diameter in each well was based on normalization to the amount of myotubes quantified. Only myotubes that were entirely visible within each field of view were eligible for analyses. All analysis of myotube size was performed by the same, blinded investigator using ImageJ ^41^.

### Statistics

Data were collected using Excel 2016 and analyzed with GraphPad Prism 10.1.1 (GraphPad Software), R Studio (v4.5.0), and STRING v12. Data are expressed as mean ± SD, including individual values where applicable. Significance was set at α=0.05, with p<0.05 shown. Data distribution was assessed using Shapiro–Wilk tests. Ordinary one-way ANOVA with Tukey’s post-hoc test was used for comparing one variable across more than two groups. Two-way ANOVA with Tukey’s post-hoc test was used for comparing multiple variables across both groups. Student’s t-test or Mann-Whitney tests were used to compare one variable between both groups. Fiber CSA frequency distributions were analyzed using linear mixed-effects models followed by bin-wise post hoc comparisons corrected using the Benjamini-Hochberg false discovery rate. Log2FC comparisons between groups were made using a Welch’s t-test while permutation was applied to adjust for multiple comparisons. Data plots were generated with GraphPad Prism and R Studio. Unless otherwise stated, analyses were performed using all available samples for each assay. Missing assay-level data were not imputed, and sample numbers therefore varied according to sample/material availability and technical feasibility.

### Study approval

The study was conducted in accordance with the Declaration of Helsinki, approved by the Regional Scientific Ethical Committee of the Capital Region of Denmark (H-21035808), and registered at ClinicalTrials.gov (NCT05307367). Written informed consent was obtained from all participants before study participation. Approval for processing of clinical data was granted by the local institutional board (514-0685/22-3000). Clinical data were stored on a REDCap server operated by the Capital Region of Denmark in accordance with the European Union General Data Protection Regulation (GDPR), and a data transfer agreement was obtained (Jr.nr 22019423). No patient-identifiable photographs are included in this manuscript.

### Data sharing statement

The proteomics dataset generated in this study, together with supporting analysis files, will be made publicly available at [repository name; accession/DOI to be added upon acceptance]. Additional source data underlying the figures are available from the corresponding author upon reasonable request.

### Role of the funding source

The funders had no role in study design, data collection, data analysis, data interpretation, writing of the report, or the decision to submit the manuscript for publication.

## Results

### Structural alterations of the skeletal muscle microenvironment in patients with NSCLC

To define structural remodeling of skeletal muscle in patients with NSCLC, we assessed muscle fiber cross-sectional area (CSA), lipid deposition and fibrosis in skeletal muscle biopsies from newly diagnosed NSCLC patients and weight- and sex-matched controls. Baseline participant characteristics are included in Table 1.

Mean muscle fiber CSA was markedly reduced in NSCLC patients compared to controls (Figure 1A-B). This was primarily driven by a 23.5% reduction in type II fiber CSA, whereas reductions in type I fiber CSA were less pronounced (Figure 1B). Fiber type proportions were similar between groups, indicating that the reduction in fiber size was not explained by major differences in fiber type composition (Figure 1C). Frequency distribution analysis showed altered fiber size distributions in NSCLC muscle, with a higher proportion of smaller type II fibers in NSCLC (P < 0.0001) and a similar trend in type I fibers (P = 0.0509; Figure 1D–G).

**Figure 1.**
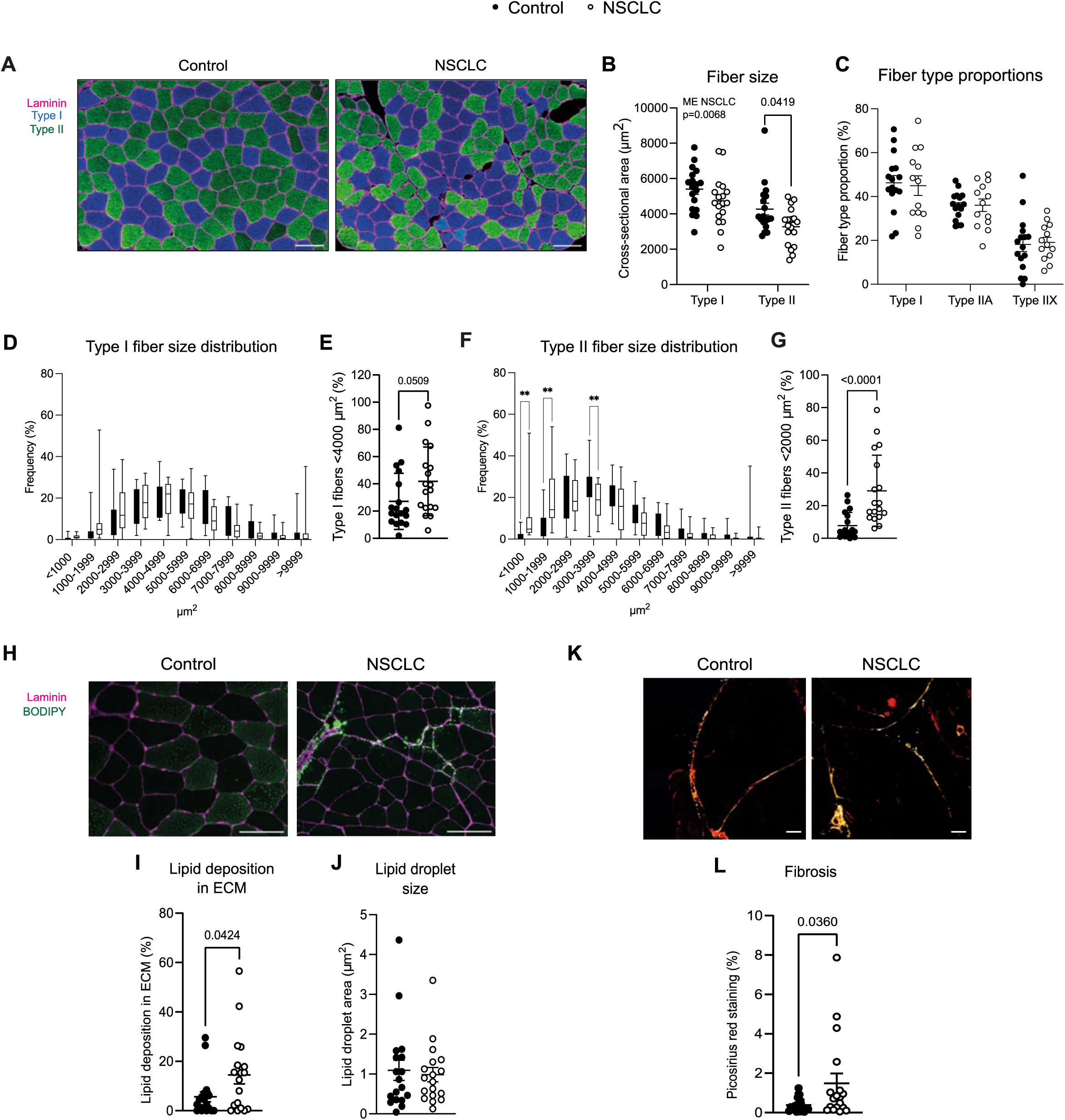
Skeletal muscle microenvironment structural remodeling in patients with NSCLC. (A) Representative cryosections of skeletal muscle from controls and patients with NSCLC stained for laminin (magenta), type I fibers (blue), and type II fibers (green). (B) Quantification of fiber cross- sectional area (CSA, µm²) in type I and type II fibers. Data points represent mean fiber CSA for each participant. (C) Fiber type proportions (%). (D) Frequency distribution of type I fiber CSA and (E) percentage of type I fibers <4000 µm². (F) Frequency distribution of type II fiber CSA and (G) percentage of type II fibers <2000 µm². (H) Representative images of BODIPY (green) and laminin (magenta) staining. Quantification of (I) lipid deposition in the extracellular matrix (ECM; % fibers) and (J) lipid droplet size (μm^2^). (K) Representative images of Picrosirius red staining and (L) quantification of fibrosis (% Picrosirius red-positive area). Statistical analyses: (B,C) two-way ANOVA; (D,F) linear mixed-effects models with Benjamini-Hochberg-corrected post hoc comparisons; (E,G,I,J,L) Student’s t-test. ME, main effect. Scale bars: (A,H,K) 100 µm.

Because chronic muscle wasting conditions, such as muscular dystrophies and sarcopenia, are often associated with fibro-fatty ECM remodeling^47^, we next evaluated muscle lipid distribution and fibrosis. BODIPY staining showed that patients with NSCLC had a redistribution of muscle lipids to the ECM (Figure 1H-I, Supplementary Figure 1C). In contrast, lipid droplet area was similar between groups (Figure 1J), while overall intramyocellular lipid intensity was lower in NSCLC muscle across both fiber types (Supplementary Figure 1D-E). Additionally, we observed a trend towards increased type I/type II intensity ratio in NSCLC, suggesting relatively greater lipid retention in oxidative fibers (Supplementary Figure 1F). Despite these alterations in lipid distribution, intramuscular glycogen deposition was similar between groups (Supplementary Figure 1G). In parallel, collagen deposition was greater in patients with NSCLC compared with controls (Figure 1K-L), indicating fibrotic remodeling alongside extracellular lipid redistribution.

Interestingly, the muscle protein content of the atrophy-related genes (atrogenes) ATROGIN1 and MuRF1, together with the autophagy marker P62, were similar between patients with NSCLC and controls (Supplementary Figure 1H-J), in line with findings from patients with gastric cancer^48^.

Together, these data show structural remodeling in NSCLC muscle characterized by preferential atrophy of type II fibers, extracellular lipid redistribution, and fibrosis. These structural alterations prompted us to next examine global proteomic remodeling in NSCLC skeletal muscle.

#### Skeletal muscle pathological proteomic remodeling in patients with NSCLC

Having established structural remodeling of the muscle microenvironment in NSCLC, we next employed unbiased mass spectrometry-based global proteomics to define molecular alterations in skeletal muscle from patients with NSCLC compared with controls. Principal component analysis showed clear separation between groups (Figure 2A). In total, 3597 proteins were quantified in the dataset, of which 905 were increased and 541 decreased in NSCLC skeletal muscle compared with controls (adjusted *P* < 0.05; Figure 2B).

**Figure 2.**
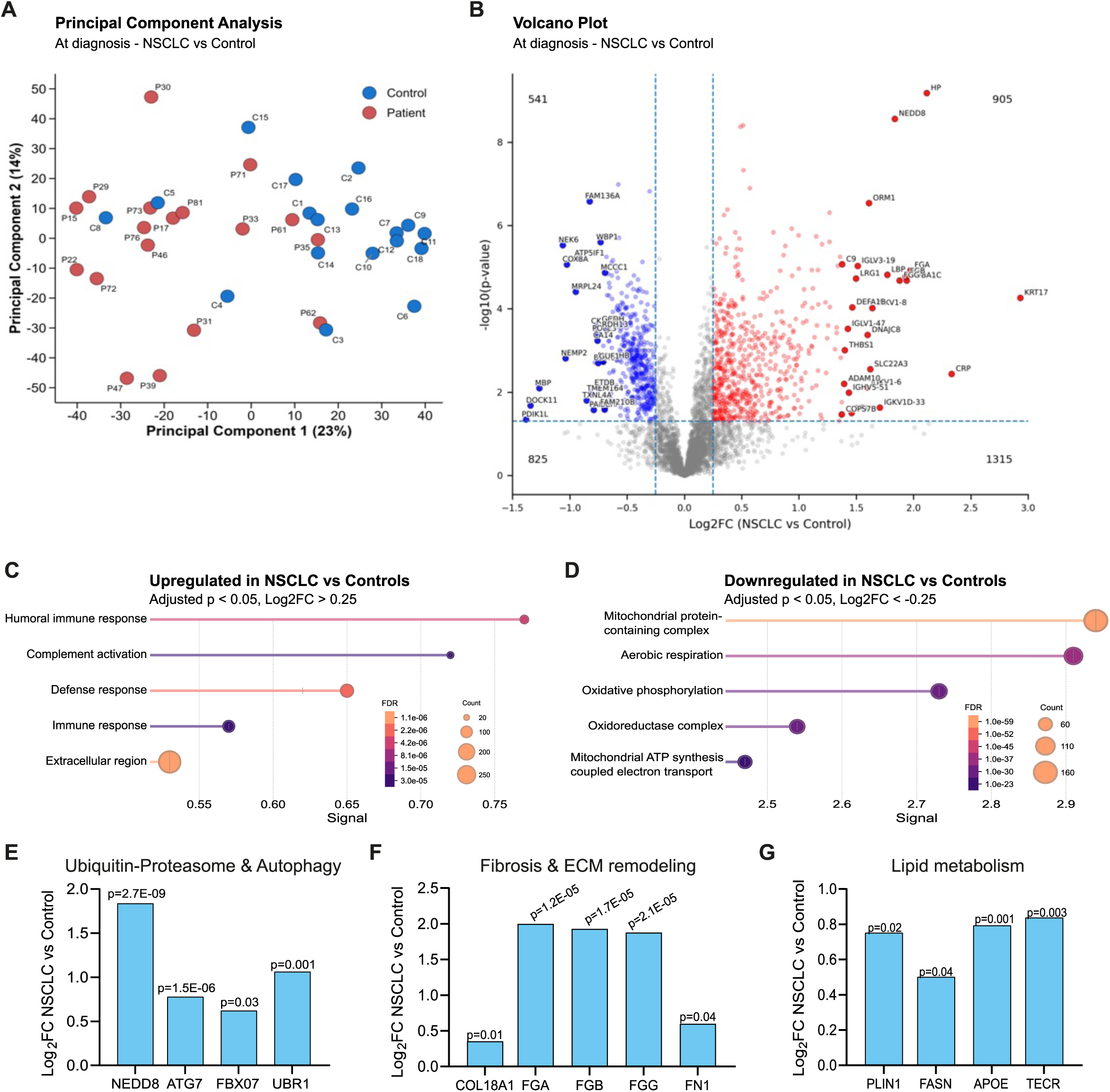
Skeletal muscle proteomic remodeling in patients with NSCLC. (A) Principal component analysis of skeletal muscle proteomes from controls and patients with NSCLC. (B) Volcano plot showing log₂ fold change (log₂FC; NSCLC vs. control) and −log₁₀ P values for quantified proteins. (C,D) Gene Ontology over-representation analysis of proteins increased (C) or decreased (D) in NSCLC skeletal muscle. (E–G) Differential abundance of selected proteins involved in ubiquitin-proteasome/autophagy pathways (E), fibrosis and extracellular matrix (ECM) remodeling (F), and lipid metabolism (G). Statistical analysis: (E–G) Welch’s t-test.

To identify broader biological patterns associated with these changes, we performed enrichment analysis on proteins increased and decreased in NSCLC muscle (adjusted *P* < 0.05, Log2FC 0.25). Among proteins increased in NSCLC, there was an over-representation of humoral immune response, complement activation, defense response, immune response, and extracellular region terms (Figure 2C). In contrast, amongst proteins decreased in NSCLC muscle, there was an over-representation of mitochondrial and bioenergetic terms, including mitochondrial protein-containing complex, aerobic respiration, oxidative phosphorylation, oxidoreductase complex, and mitochondrial ATP synthesis coupled electron transport (Figure 2D). Ranked gene set enrichment analysis supported this overall pattern, confirming enrichment of immune/extracellular terms among proteins increased in NSCLC and mitochondrial/bioenergetic terms among proteins decreased in NSCLC (Supplementary Figure 2A).

In addition to these global enrichment patterns, inspection of selected differentially abundant proteins highlighted pathways consistent with the structural changes observed histologically (Figure 1). Proteins linked to ubiquitin-proteasome function and autophagy, including NEDD8, ATG7, FBXO7, and UBR1, were increased in NSCLC muscle (Figure 2E). Likewise, proteins related to fibrosis and ECM remodeling, including COL18A1, FN1, FGA, FGB, and FGG, were increased (Figure 2F). Selected enrichment analysis also identified lipid-binding and lipid-transport pathways, in line with the histological findings of extracellular lipid redistribution (Supplementary Figure 2B). In agreement with this, several proteins involved in lipid handling and metabolism, including PLIN1, FASN, APOE, and TECR, were increased in NSCLC muscle (Figure 2G).

Overall, these data indicate broad proteomic remodeling in NSCLC skeletal muscle, characterized by an over-representation of inflammatory and ECM terms amongst upregulated proteins, and mitochondrial terms amongst downregulated proteins. These observations guided the subsequent analyses of inflammatory and immune-cell remodeling, mitochondrial and calcium-handling alterations, and stromal/myogenic niche changes in NSCLC muscle.

#### Enhanced inflammatory and immune responses in NSCLC skeletal muscle

Proteomic profiling identified increased inflammatory and extracellular programs in NSCLC skeletal muscle, including enrichment of immune response, humoral immune response, complement activation, and defense response terms (Figure 2C). In line with this, several inflammatory and acute-phase proteins were increased in NSCLC muscle, including CRP, HP, ORM1, S100A8, and S100A9, together with the complement component C3 (Figure 3A). Notably, C3 has recently been implicated as a potential mediator of inflammation-associated muscle atrophy in pancreatic cancer ^49^, suggesting that complement-related signaling may contribute to muscle remodeling across cancer types. These findings prompted us to investigate inflammatory signaling and immune cell composition in the NSCLC muscle microenvironment in more detail.

**Figure 3.**
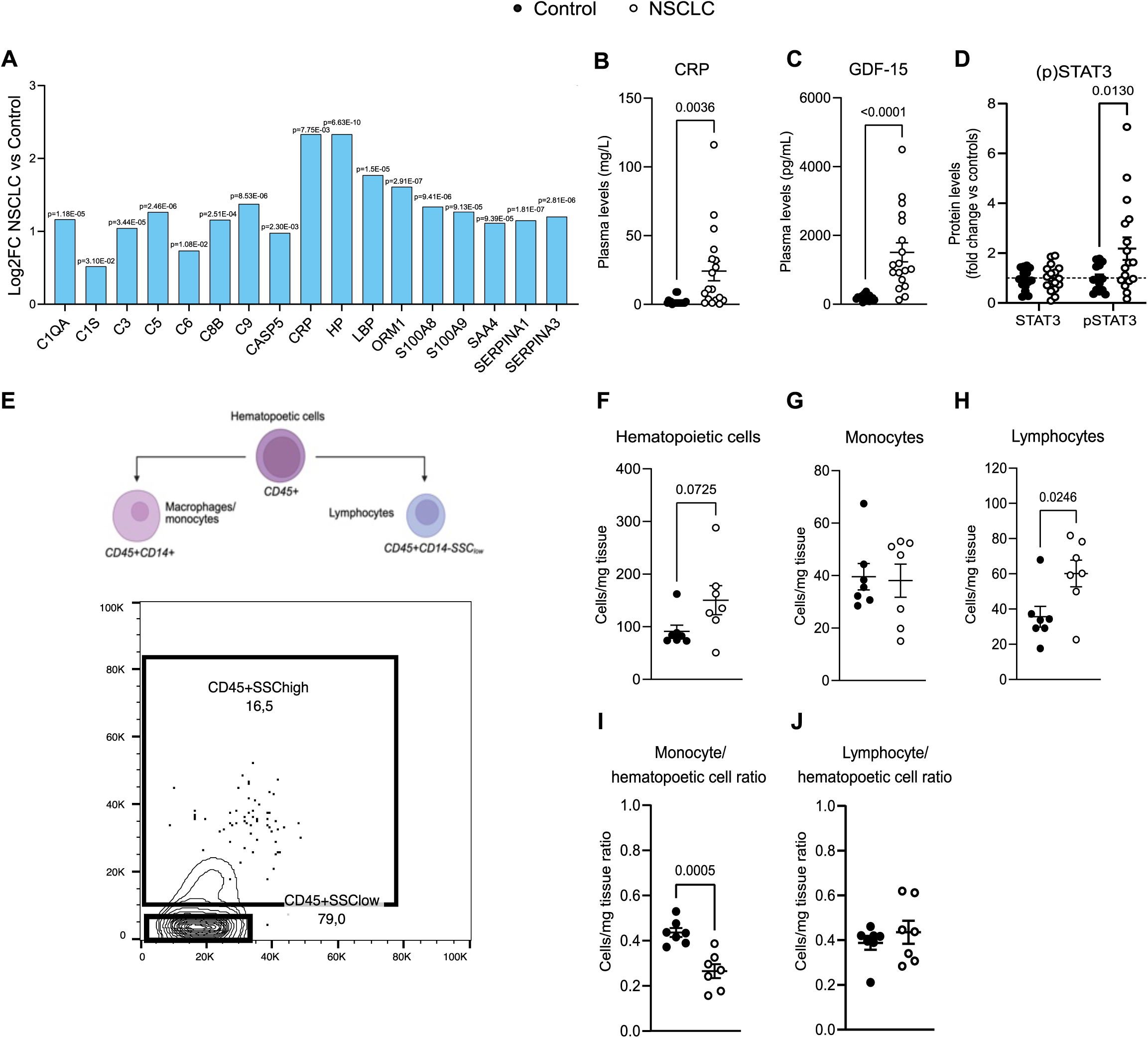
Inflammatory and immune remodeling in NSCLC skeletal muscle. (A) Log₂ fold changes (log₂FC; NSCLC vs. control) in selected inflammatory, acute-phase, complement, and cytokine- associated proteins identified by proteomics. (B,C) Plasma CRP (B) and GDF-15 (C) concentrations. (D) Quantification of STAT3 and phosphorylated STAT3 (pSTAT3) protein abundance. (E) Summary of the flow cytometry gating strategy used to identify immune-cell subsets and flow plot with the CD45^+^CD14^-^ population divided into lymphocytes (side scatter low) and granulocytes (side scatter high). (F) Hematopoietic cell content (CD45⁺; cells per mg tissue). (G) Monocyte/macrophage content (CD45⁺CD14⁺; cells per mg tissue). (H) Lymphocyte content (CD45⁺CD14^−^SSC^low^; cells per mg tissue). (I) Monocyte/macrophage-to-hematopoietic cell ratio and (J) lymphocyte-to-hematopoietic cell ratio. Statistical analyses: (B,C,F–J) Student’s t-test; (D) two-way ANOVA.

At the systemic level, patients with NSCLC had increased plasma CRP and GDF-15 concentrations compared with controls (Figure 3B-C). In contrast, GDF-15 mRNA expression in skeletal muscle showed only a trend towards an increase in NSCLC (Supplementary Figure 3A), and analysis of inflammatory gene expression in muscle revealed substantial heterogeneity within the NSCLC group (Supplementary Figure 3B). In three patients, *IL6* gene expression was approximately 2.5-fold, 7.4-fold and 40-fold greater than controls (Supplementary Figure 3B). To evaluate a potential activation of IL6 signaling in skeletal muscle in the context of NSCLC, we quantified protein content of its downstream effectors NF-κB subunit P65 and STAT3. Whereas total STAT3 and P65 protein levels were similar between groups, pSTAT3 was increased in NSCLC muscle (Figure 3D, Supplementary Figure 3C), consistent with activation of STAT3-associated inflammatory signaling.

We next assessed whether this inflammatory profile was accompanied by altered immune cell composition using flow cytometry (Figure 3E). This analysis identified a trend towards an increased infiltration of CD45^+^ hematopoietic cells (Figure 3F). Within this compartment, total monocyte content was similar between groups (Figure 3G), whereas lymphocyte content was increased in NSCLC muscle (Figure 3H). Within the myeloid compartment (CD45^+^CD14^+^), we separated macrophages/monocytes according to their inflammatory profile using CD11c ^50^ (Supplementary Figure 3E-F), revealing no differences in macrophage subsets between groups. Relative quantification further showed a reduced monocyte-to-hematopoietic cell ratio in NSCLC compared with controls (Figure 3I), while the lymphocyte-to-hematopoietic cell ratio was unchanged (Figure 3J). These findings indicate a lymphoid bias within NSCLC skeletal muscle.

Together, these findings indicate inflammatory and immune remodeling in NSCLC muscle, characterized by increased inflammatory and complement-associated proteins, elevated circulating CRP and GDF-15, increased pSTAT3, and altered immune-cell composition with greater lymphocyte abundance and a reduced relative monocyte representation. This altered immune profile and STAT3 activation aligns with previous studies showing that immune cell infiltration in patients with cancer correlates with decreased muscle mass and altered muscle morphology ^10,11^.

#### NSCLC skeletal muscle exhibits mitochondrial alterations, oxidative stress and calcium dysregulation

Because our proteomic analysis identified coordinated downregulation of bioenergetic pathways in NSCLC skeletal muscle, we proceeded with examination of features related to mitochondrial biology. Further exploration of the proteomics dataset revealed reduced abundance of proteins linked to mitochondrial respiration, translation, redox balance, and membrane organization, including COX8A, MRPL24, NNT, OXA1L, TXN2, and PRDX3, in NSCLC muscle (Figure 4A).

**Figure 4.**
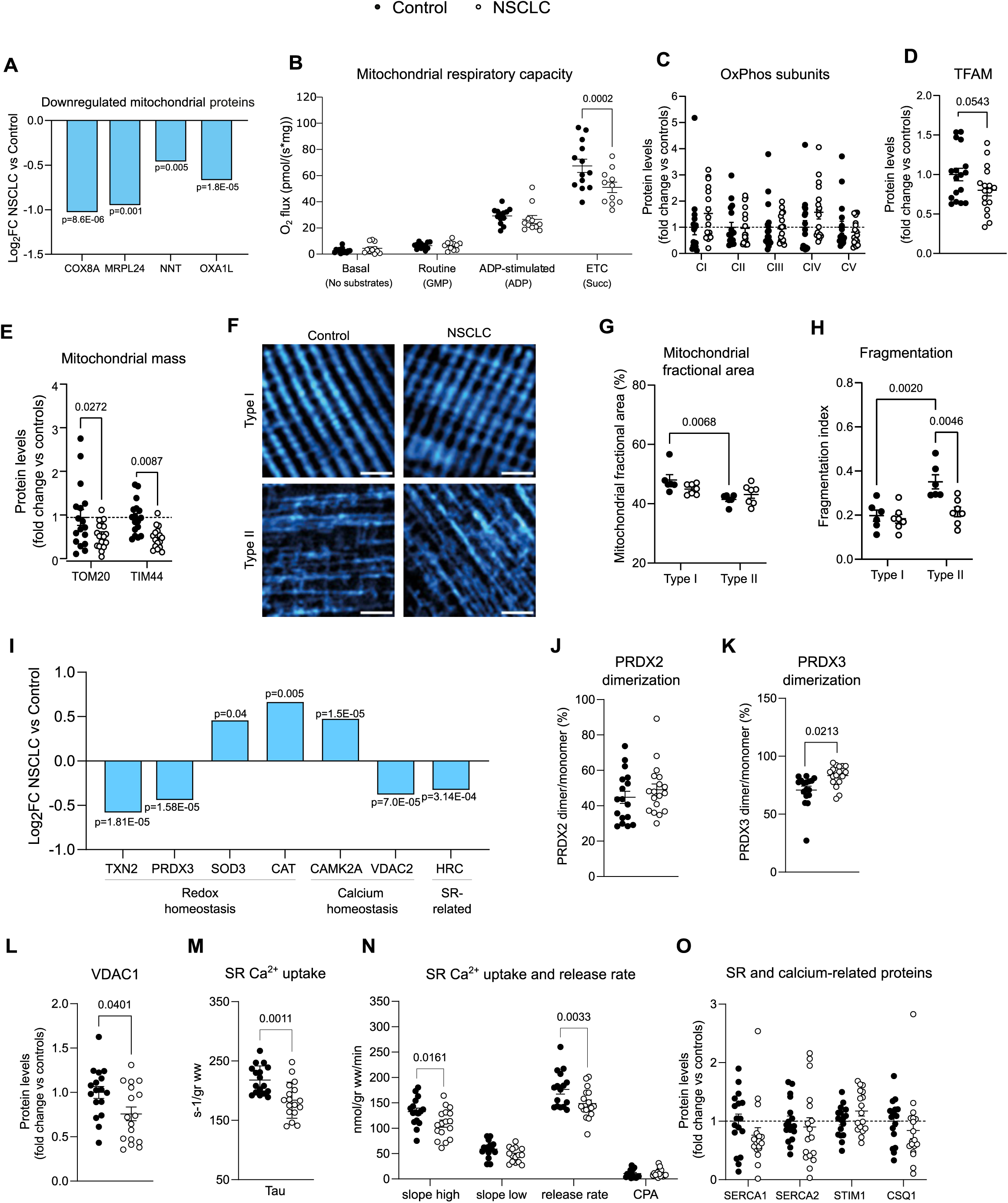
Mitochondrial and calcium-handling remodeling in NSCLC skeletal muscle. (A) Log₂ fold changes (log₂FC; NSCLC vs. control) in selected downregulated mitochondrial proteins identified by proteomics. (B) Mitochondrial respiratory capacity measured as oxygen flux per muscle mass. (C-E) Quantification of OXPHOS subunits (C), TFAM protein abundance (D), and mitochondrial mass markers TOM20 and TIM44 (E). (F) Representative immunofluorescence images of type I and type II muscle fibers stained for COXIV to visualize mitochondrial network organization (cyan). Quantification of mitochondrial fractional area (G) and fragmentation index (H). (I) Log₂ fold changes (log₂FC; NSCLC vs. control) in selected redox-, mitochondrial-, and calcium-handling-related proteins identified by proteomics. (J,K) PRDX2 (J) and PRDX3 (K) dimer/monomer ratio. (L) VDAC1 protein abundance. (M) Sarcoplasmic reticulum (SR) Ca²⁺ uptake rate, expressed as tau. (N) SR Ca²⁺ uptake at high and low Ca²⁺ concentrations, SR Ca²⁺ release rate, and vesicle leak estimated using cyclopiazonic acid (CPA). (O) Quantification of SR and calcium-related proteins SERCA1, SERCA2, STIM1, and CSQ1. Statistical analyses: (B,C,E,G,H,N,O) two-way ANOVA; (D,J–M) Student’s t-test. Scale bar: (F) 5 µm.

To understand the functional consequences of these proteomic changes, we evaluated skeletal muscle mitochondrial oxygen consumption. We detected decreased maximal coupled oxygen consumption upon addition of succinate, along with a trend towards decreased ADP-stimulated oxygen consumption in patients with NSCLC, demonstrating reduced mitochondrial respiratory capacity (Figure 4B). These alterations were not associated with differences in the protein levels of OXPHOS subunit forming complexes (Figure 4C, Supplementary Figure 4A) nor mitochondrial transcripts (Supplementary Figure 4B). However, we detected a trend towards a decrease in protein content of the mitochondrial transcription factor TFAM (Figure 4D, Supplementary Figure 4C), which can impact mitochondrial DNA (mtDNA) stability and abundance, leading to alterations in OXPHOS subunit content and respiratory capacity.

Because respiratory capacity is influenced by mitochondrial abundance, we evaluated this parameter using several approaches. Gene expression analysis of the mitochondrial biogenesis marker PGC1α (*PPARGC1a*) showed similar levels in skeletal muscle of patients with NSCLC compared to controls (Supplementary Figure 4D). In contrast, we observed a reduction in protein levels of the mitochondrial membrane proteins TOM20 and TIM44 in skeletal muscles of patients with NSCLC compared to controls (Figure 4E, Supplementary Figure 4E), which is indicative of decreased mitochondrial mass^51^. To corroborate these findings, we used confocal imaging to quantify the mitochondrial fractional area as an estimate of mitochondrial mass. We observed lower mitochondrial fractional area in type II fibers compared to type I fibers in controls, whereas this difference was blunted in NSCLC (Figure 4F-G). Together, these findings support remodeling of mitochondrial content and distribution in NSCLC skeletal muscle.

Control of mitochondrial morphology is a crucial determinant of mitochondrial function and prevention of muscle inflammation and atrophy^52,53^. Although we did not observe differences in protein content of mitochondrial fusion proteins Mitofusin 1 and 2 (MFN1, 2) and the master regulator of fission DRP1 (Supplementary Figure 4F-G), we detected reduced fragmentation in type II fibers in NSCLC compared to controls (Figure 4H), indicating altered mitochondrial morphology in the fiber type most affected by atrophy, which may impact the oxidative capacity of muscle fibers.

Enhanced oxidative stress is tightly linked to the development of muscle atrophy ^54^. Given that our proteomic analysis showed reduced PRDX3 in patients with NSCLC (Figure 4I), we evaluated intracellular compartmentalization of oxidative stress by quantifying the dimerization ratio of peroxiredoxins (PRDX2 and PRDX3), which reside in the cytosol and mitochondria, respectively. Under high oxidative stress, PRDXs oxidize and dimerize (Supplementary Figure 4H). Our data show that PRDX2 total levels and dimerization remained unaltered (Supplementary Figure 4I-J). However, PRDX3 total levels were decreased in NSCLC compared to controls (Supplementary Figure 4I, K), consistent with our proteomic findings. Concomitantly, we observed unaltered PRDX2 dimerization ratio in NSCLC (Figure 4J), and yet, an increased PRDX3 dimerization ratio (Figure 4K), indicating elevated oxidative stress limited to mitochondria. Interestingly, the increased PRDX3 dimerization was not explained by enhanced mitochondrial superoxide dismutase (SOD2) protein content (Supplementary Figure 4I, L), which could lead to increased mitochondrial H2O2. Furthermore, we found no change in total H₂O₂ levels in skeletal muscle (Supplementary Figure 4M). These data suggest the potential contribution of other mitochondrial redox species to the dimerization of PRDX3.

In skeletal muscle, the SR regulates calcium release and uptake during muscle contraction and relaxation. Having identified alterations in calcium handling and SR-related proteins in our proteomic and western blotting analysis (Figure 4I, L), we next assessed SR function in relation to calcium handling. We found a slower SR calcium uptake and reduced SR calcium release rate in patients with NSCLC compared with controls (Figure 4M-N), despite comparable protein levels of components of SR calcium cycling machinery (SERCA 1/2, STIM1 or CSQ1; Figure 4O, Supplementary Figure 4N). These findings suggest that the functional capacity or activity of these proteins, rather than their expression levels, may be altered. Overall, these findings demonstrate impairments in the activity of SR and alterations in proteins related to calcium handling in skeletal muscle in patients with NSCLC, which is in line with our previous observations in a murine model of cancer-associated muscle wasting^55^.

Together, these findings indicate that NSCLC skeletal muscle undergoes coordinated mitochondrial and calcium-handling remodeling, characterized by altered respiratory capacity, reduced abundance of mitochondrial proteins, fiber type-dependent changes in mitochondrial morphology and fractional area, altered mitochondrial redox status, and impaired SR calcium uptake and release. Overall, these data support broad remodeling of mitochondrial and calcium-regulatory systems in NSCLC skeletal muscle, which may contribute to the development of muscle atrophy in patients with NSCLC.

#### Skeletal muscle of patients with NSCLC has a greater proportion of CD90^-^ FAPs, mechanistically driving muscle atrophy *in vitro*

Successful regeneration, adaptation, and homeostatic maintenance of skeletal muscle is largely facilitated by the coordinated interplay between tissue-resident MuSCs, FAPs and various immune cells. Furthermore, FAPs have recently emerged as crucial regulators of both ECM and muscle fiber remodeling in the context of health and disease. Therefore, in a subgroup analysis (n=7) in NSCLC and controls, respectively, we quantified MuSC and FAP content, using FACS, while isolating the cells for further downstream functional analysis ex vivo.

Overall MuSC content was similar between patients with NSCLC and controls (Figure 5A). However, the flow-mediated cell size of MuSCs was significantly increased in NSCLC muscle compared with controls (Figure 5B). While MuSC morphology and size is tightly coupled to rate of activation, we observed a marked individual variation in proliferative capacity amongst NSCLC-derived MuSCs compared to matched controls (Figure 5C-D). The discrepancy between cell morphology and ex vivo function, could suggest manifestations of increased cellular stress or senescence in MuSCs affected be NSCLC.

**Figure 5.**
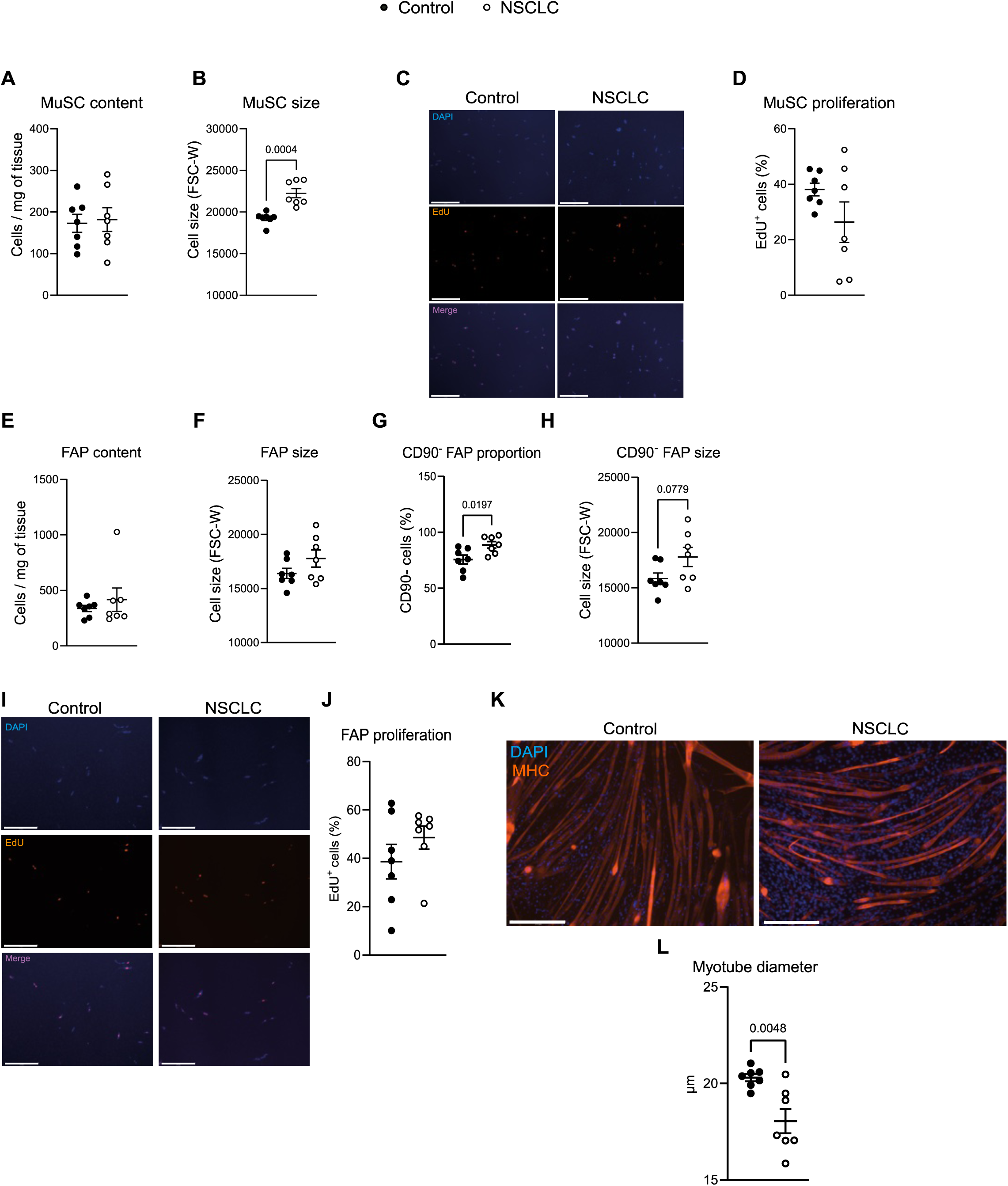
Increased CD90^−^ FAP proportion in NSCLC skeletal muscle and reduced myotube size after exposure to NSCLC-derived FAP conditioned media. (A) Muscle stem cell (MuSC; CD82⁺CD56⁺CD45^−^CD31^−^CD34^−^) content (cells per mg tissue) and (B) MuSC size (FSC-W) in controls and patients with NSCLC. (C) Representative images of MuSCs stained with DAPI (blue) and EdU, incorporated into newly synthesized DNA (orange), and (D) quantification of EdU incorporation (%) after 3 days in culture. (E) Total FAP content (cells per mg tissue) and (F) total FAP size (FSC-W). (G) CD90^−^ FAP proportion (%) and (H) CD90^−^ FAP size (FSC-W). (I) Representative images of FAPs stained with DAPI (blue) and EdU (orange), and (J) quantification of EdU incorporation (%) after 5 days in culture. (K) Representative images of C2C12 myotubes treated with conditioned media from control- or NSCLC-derived FAPs and (L) quantification of myotube diameter. Statistical analyses: (A,B,D–H,J,L) Student’s t-test. Scale bars: (C,I,K) 275 µm.

As in the case for MuSCs, overall FAP content was similar between patients with NSCLC and controls (Figure 5E). For overall FAPs, we did not find significant differences in cell size between patients and controls (Figure 5F). However, FAPs are characterized by a remarkable cellular heterogeneity, with functionally distinct subsets displaying different molecular signatures. Therefore, we discriminated FAPs according to CD90 expression, based on our previous work ^43^. This revealed a increased proportion of CD90^-^ FAPs in NSCLC compared with controls (Figure 5G). In addition, a trend for increased cell size of the CD90^-^ FAPs was observed in NSCLC-patients compared to controls (Figure 5H), which could indicate increased metabolic activity of these cells, specifically, in NSCLC. Interestingly, CD90^-^ FAPs have previously been suggested to hold a greater adipogenic differentiation potential compared to CD90^+^ FAPs. Analysis of FAP proliferation capacity indicated similar propensity of NSCLC patient-derived FAPs to enter the cell cycle compared to controls (Figure 5I-J).

Emerging evidence identifies FAPs as important secretory cells, mechanistically providing direct trophic support for muscle fiber maintenance and growth^28,30,56,57^. Therefore, we performed conditioned media experiments to investigate the mechanisms of action of FAP secreted factors on C2C12 myotube growth (Supplementary Figure 5). Here, we found that media conditioned with patient-derived FAPs markedly reduced myotube width compared to media conditioned with control-derived FAPs (Figure 5K-L), demonstrating a mechanism whereby altered paracrine signaling from the muscle-resident stromal compartment directly drives cachexia-associated muscle wasting. In relation to the increased markers of inflammation and C3 expression in NSCLC patients, we have recently identified FAPs as a highly specific local source of C3 in both young and elderly human skeletal muscle ^32^. Given our observations of increased CD90^-^ FAP predominance in NSCLC skeletal muscle (Figure 5G), we evaluated whether these differences in FAP subsets could explain the increased C3. In support of this hypothesis, single-cell RNA sequencing data confirmed the enrichment of C3 RNA expression in CD90^-^ FAPs compared to CD90^+^ FAPs (Supplementary Figure 5B), thus demonstrating a potential link between FAP subpopulation shifts and altered inflammatory signaling in NSCLC.

Together, these findings indicate that the composition of FAPs within the skeletal muscle microenvironment is altered in the context of NSCLC, and that these alterations are associated with a functional switch of the tissue-resident FAP population that may mechanistically contribute to muscle atrophy.

## Discussion

Here, we comprehensively characterize skeletal muscle and its microenvironment in patients with advanced stage NSCLC, providing clinically relevant insights into proteomic rewiring and molecular alterations that mechanistically could underlie the high propensity for cancer-induced muscle wasting in this population ^5^. We reveal a multifaceted signature of cancer- associated muscle remodeling, encompassing fibrosis, lipid redistribution, mitochondrial adaptations, immune cell reorganization, and altered FAP populations, which mechanistically links paracrine signaling from the muscle-resident stromal compartment to atrophy in cancer cachexia.. Together, these findings establish a unique human proteomic resource and highlight potential mechanisms contributing to muscle wasting and inflammation.

We found that NSCLC is associated with skeletal muscle wasting driven primarily by type II muscle fiber atrophy. This aligns with observations in rodent models of cancer-induced muscle wasting^58,59^. In patients with various cancer types, muscle atrophy was observed in both type I and type II chemically skinned muscle fibers^60^. Thus, fiber-type selective atrophy may be cancer-type specific. The preferential type II muscle fiber atrophy in NSCLC may be explained by greater susceptibility to proteolytic pathways^10,11,61^. Our findings indicate that this phenomenon may be attributable to altered mitochondrial homeostasis, as evidenced by fiber type II-specific adaptations in mitochondrial network organization in patients with NSCLC.

In addition to muscle atrophy, we observed increased fibrosis and a redistribution of lipids toward the ECM in patients with NSCLC. Fibrosis has been implicated in muscle wasting in both patients with pancreatic cancer and preclinical models of lung cancer^23,24^. The extracellular lipid accumulation we report builds upon previous observations that cancer is associated with reduced skeletal muscle density and fat infiltration into muscle tissue^26,27,62^. While earlier studies reported increased lipid droplet content or overall muscle fat infiltration, our results demonstrate a reduction in total intramyocellular lipid content. This discrepancy may reflect methodological differences, as we directly quantified lipid droplets via immunohistochemistry, whereas prior studies relied on CT-derived radiodensity^26,62^ or electron microscopy^27^. Yet, our findings indicate that in NSCLC, lipids are redistributed toward the ECM, pointing to a potential fibro-adipogenic origin for this shift.

Our findings support a model in which the fibro-fatty remodeling observed in patients with NSCLC plays a role in the development of skeletal muscle atrophy and dysfunction^63^. While overall FAP content was similar between groups, the shift towards CD90^-^ FAPs in patients with NSCLC could partly explain their predisposition towards ECM fat deposition. We propose that the lipid deposition in NSCLC is mechanistically driven by this enrichment of pro-adipogenic CD90^-^ FAPs, as specific FAP subpopulations have been shown to contribute to fatty infiltration in muscle degeneration.

Previous pre-clinical studies have shown that FAP-derived factors crucially impact muscle fiber homeostasis, partly through modulation of the local muscle environment^56,57^. Importantly, our findings uncover a mechanism by which conditioned media from patient-derived FAPs induces myotube atrophy *in vitro*, implicating FAPs as a local driver of muscle atrophy upon cancer. The ability of FAPs to reduce myotube size independently of direct immune or tumor cell interaction suggests that they may autonomously induce muscle wasting in NSCLC in a paracrine manner. Interestingly, these findings are in line with previous observations in preclinical models of muscle wasting^64^ and muscle denervation^30^. To our knowledge, our findings are the first to identify direct FAP-mediated loss of myogenic support in the context of cancer-associated muscle atrophy in humans. These findings strongly warrant further investigation of FAP biology in the context of chronic disease, and advocates for targeting FAPs or their mechanistic role as paracrine mediators to attenuate muscle wasting in NSCLC.

Together with a greater proportion of CD90^−^ FAPs, we also observed immune cell infiltration in the skeletal muscle of patients with NSCLC, which aligns with emerging insights into FAP-mediated immune modulation during muscle regeneration. Pre-clinical studies have revealed cross-talk from FAPs to both macrophages^65,66^ and muscle-resident T-cells^67^. Recently, it has been demonstrated in aged human skeletal muscle that FAPs are a local source of C3, a complement factor that supports recruitment, survival, and phagocytic activity of pro-inflammatory macrophages during regeneration^32^. Our proteomic analyses revealed increased abundance of C3, together with other pro-inflammatory markers, such as CRP, HP, S100A8 and S100A9, in the skeletal muscle of patients with NSCLC. This elevation of C3 in NSCLC aligns with a recent report highlighting complement pathway activation in skeletal muscle from pancreatic cancer-induced muscle wasting^49^. Notably, our sequencing data from human skeletal muscle clearly demonstrates enrichment of C3 transcripts in CD90^-^ FAPs compared to CD90^+^ FAPs. Recent work identifying molecular subtypes in human skeletal muscle upon cancer provides strong support for our findings^68^. In that study, transcriptome analysis revealed that patients in the cachexia-associated subtype had higher expression of immune/cytokine storm pathways and ECM remodeling, with type IIA and IIX fiber atrophy identified through histological analysis. Our study builds on and complements these findings by providing the first proteomic evidence of increased skeletal muscle abundance of complement C3 and inflammatory mediators such as CRP, HP, S100A8 and S100A9 in patients with NSCLC. Together, these complementary transcriptomic and proteomic data provide converging evidence that inflammation and complement activation are central features of the cachectic muscle phenotype, reinforcing their potential as therapeutic targets.

We also observed elevated circulating levels of GDF-15, which has been implicated in the anorexia component to cancer-driven muscle wasting^12^, and more recently in tumor-associated immunity ^69,70^, while its effects on the muscle remain unclear. Concomitantly, we detected STAT3 activation, a major intrinsic signal responsive to a wide range of cytoplasmic receptors, leading to immune modulatory pathways^71^. These data highlight the pro-inflammatory milieu of skeletal muscles of patients with NSCLC, which can further contribute to the development of muscle atrophy. Importantly, oxidative stress induced by H2O2 treatment has been reported to STAT3 phosphorylation in human lymphocytes^72^. STAT3 is phosphorylated and translocated onto mitochondria upon exacerbated mitochondrial ROS production induced by Complex I inhibition in human cultured cells^73^. In this regard, we show that while total H2O2 levels remained unchanged, the selective increase in PRDX3 dimerization suggests enhanced oxidative stress restricted to mitochondria in patients with NSCLC, where alternative redox species to H2O2 may be involved. Furthermore, a study using a murine model for denervation-induced muscle atrophy demonstrates that skeletal muscle ROS activates neutrophils and attracts them to the tissue^74^, highlighting a direct link between skeletal muscle oxidative stress and immune cell infiltration. Given these reported studies and our observations, we propose that FAP redistribution, enhanced C3 production and oxidative stress in NSCLC muscles synergistically promote immune cell infiltration and further muscle atrophy in this context.

Mitochondrial abnormalities have been implicated in cancer-driven muscle atrophy, with preclinical and human studies suggesting that altered mitochondrial mass, distribution and respiration exacerbate muscle wasting^16,17,75,76^. Our data show that patients with NSCLC exhibited decreased respiratory capacity and mass, and type II fiber-specific reductions in mitochondrial fragmentation. Notably, it has been previously reported that downregulation of TOM20 in a human cancer cell line ^77^ or TIM44 in isolated mouse cardiomyocytes ^78^ result in impaired mitochondrial oxygen consumption and increased ROS production, therefore providing a potential direct molecular link between TOM20 and TIM44 reduced expression and the mitochondrial phenotype observed in patients with NSCLC.

Altogether, our data provide a comprehensive characterization of mitochondrial parameters in skeletal muscles of patients with NSCLC, and corroborate the notion that alterations in mitochondrial homeostasis are a key hallmark of cancer-related pathological skeletal muscle remodeling. Importantly, an emerging field of study describes that primary mitochondrial dysfunctions can activate inflammatory signals^18,79^. In skeletal muscle, such mechanisms result in muscle atrophy and subsequent immune cell infiltration ^52,53,80,81^. Although immune cells may exert both protective and deleterious effects in muscle ^10,11,74^, we speculate that chronic mitochondrial adaptations and oxidative stress induced by NSCLC facilitate the deleterious effects of chronic immune cell infiltration in the muscle, exacerbating the muscle inflammatory and wasting phenotype. Given that mitochondrial dysfunction and inflammation form a vicious cycle in cancer-induced muscle wasting^79^, targeting oxidative stress may represent a promising intervention to mitigate muscle loss in NSCLC.

Our data indicate disrupted calcium homeostasis at the SR and mitochondria in patients with NSCLC. We observed downregulation of the mitochondrial calcium transporter VDAC1, which may limit mitochondrial calcium buffering capacity, contributing to chronically elevated cytosolic calcium levels and stressing mitochondrial function^82–84^. According to our proteomics data, CAMK2A, a kinase implicated in denervation-mediated muscle atrophy^85^, was upregulated in NSCLC muscle, which may reflect a compensatory mechanism to restore calcium balance at the SR-mitochondrial interface^85,86^. Consistent with this, our functional measurements show impaired SR calcium uptake and release in NSCLC, and our analysis in a related study showed upregulation and phosphorylation of the ryanodine receptor in patients with CAC compared to patients without CAC ^36^. Interestingly, disrupted calcium handling can lead to elevated cytosolic calcium, which is known to activate catabolic pathways (such as calpains and NF-κB) and contribute to oxidative stress and inflammation^87^. Thus, these calcium disturbances, together with mitochondrial alterations and redox imbalance, could contribute to muscle atrophy and promote the inflammatory environment observed in NSCLC skeletal muscle^87^. Calcium-driven oxidative stress and inflammation may also contribute to the changes in FAPs we observed, since redox and inflammatory cues shape FAP fate and function^88^.

The cross-sectional design and assay-specific variation in sample availability should be taken into account when interpreting the findings. Nevertheless, the multimodal analysis of human skeletal muscle provides an integrated view of NSCLC-associated muscle remodeling and identifies several candidate pathways for further therapeutic investigation.

Collectively, we identified alterations in FAPs, calcium handling, oxidative stress, inflammation, and mitochondrial homeostasis in NSCLC, all of which are also modulated by exercise^89,90^. Consistent with this, structured exercise after chemotherapy has been shown to improve both survival and functional outcomes in patients with colon cancer^91^. Our findings raise the possibility that exercise may confer part of this benefit by mitigating the highlighted pathophysiological alterations in the skeletal muscle microenvironment and thereby improving outcomes in NSCLC. Taken together, these findings warrant future well-controlled intervention studies to define how the muscle microenvironment responds to exercise in individuals with cancer.

We provide novel insights into the complex interplay between cellular and molecular changes underlying muscle atrophy and structural remodeling in patients with NSCLC. Our analysis reveals shifts in FAP and immune cell subsets within the muscle microenvironment, along with mitochondrial alterations, enhanced oxidative stress and altered calcium handling. These interactions may drive muscle atrophy in patients with NSCLC and highlight potential therapeutic targets for counteracting cancer-associated muscle wasting.

## Author contributions

Conceptualization: E.B., A.I., JW., J.S, J.F., and L.S. Data curation: E.B., A.I., J.W., C.T.V., N.Ø., J.F., and L.S. Formal analysis: E.B., A.I., J.W., J.S., C.T.V., J.L.M., S.T., O.D., N.Ø., S.K., J.F., and L.S. Funding acquisition: J.S., J.F., and L.S. Investigation: E.B., A.I., J.W., J.S., C.T.V., J.L.M., S.T., C.K., N.R.A., E.F., S.K., N.H., A.M., S.H.R., C.P., S.L., O.D., N.Ø., J.F., L.S. Methodology: E.B., A.I., J.W., J.S., C.T.V., C.P., S.L., O.D., N.Ø., J.F., and L.S. Project administration: E.B., A.I., J.W., J.S., J.F. and L.S. Resources: J.S., J.F., and L.S. Software: E.B., A.I., J.W., C.T.V., O.D., N.Ø., and J.F. Supervision: J.F. and L.S. Validation: E.B., A.I., J.W., J.S., C.T.V., J.L.M., O.D., N.Ø., J.F., and L.S. Visualization: E.B., A.I., J.W., O.D., and C.T.V. Writing original draft: E.B. and A.I. Writing review & editing: all authors.

## Acknowledgements

We thank Professor Erik Richter and Betina Bolmgren for carrying out plasma GDF-15 analysis. We acknowledge the Core Facility for Integrated Microscopy, Faculty of Health and Medical Sciences, University of Copenhagen. We also thank the Proteomics Research Infrastructure (PRI), University of Copenhagen, for their support in performing the proteomic analyses for this study. Finally, we thank the FACS Core unit at Department at Biomedicine, Aarhus University for providing the necessary equipment for flow-based analyses. This work was supported by a research grant from the Danish Diabetes and Endocrine Academy (postdoctoral fellowship to E.B.), which is funded by the Novo Nordisk Foundation, grant number NNF22SA0079901. This work was funded by grants to LS from the Independent Research Fund Denmark (IRFD), Inge Lehmann Program and The Dansh Cancer Society, and by grants to JS from Sejer Persson and Lis Klüwer Perssons Legat, A.P. Møller Fonden til Lægevidenskabens Fremme and Danish Medical Society Research Foundation. SHR was funded by Independent Research Fund Denmark (2030-00007A), the Lundbeck Foundation (R380-2021-1451).

## Declaration of interests

The authors declare no competing interests.

## Declaration of generative AI and AI-assisted technologies in the manuscript preparation process

During the preparation of this work, the authors used ChatGPT (OpenAI) to assist with language editing and manuscript restructuring. After using this tool, the authors reviewed and edited the content as needed and take full responsibility for the content of the publication.

## Supplementary Information

*Pathophysiological remodelling of the skeletal muscle microenvironment in patients with lung cancer*

---

**Supplementary Figure 1.**
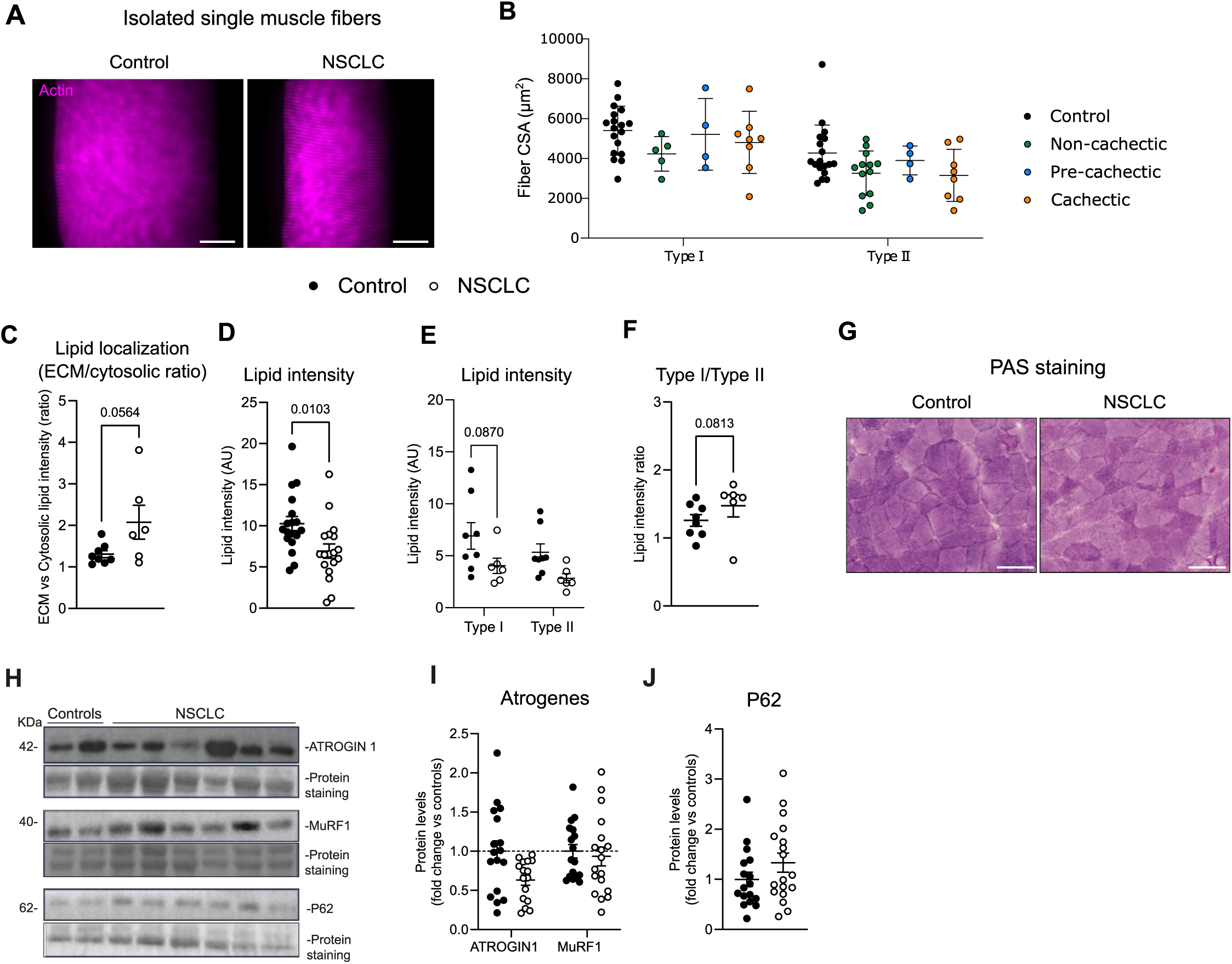
Additional histological and atrophy-related analyses in skeletal muscle from patients with NSCLC. (A) Representative images of isolated single muscle fibers stained for actin. (B) Fiber CSA stratified by clinically defined cachexia status. (C) Lipid droplet area. (D) Total lipid intensity. (E) Fiber type-specific lipid intensity in type I and type II fibers. (F) Type I/type II lipid intensity ratio. (G) Representative PAS staining of skeletal muscle from controls and patients with NSCLC. (H) Representative immunoblots of ATROGIN1, MuRF1, and P62, with total protein staining used as loading control. Quantification of (I) ATROGIN1 and MuRF1 protein abundance and (J) P62 protein abundance. (B,C,H) Two-way ANOVA. (D,I) Student’s t-test. Scale bars: 100 μm.

**Supplementary Figure 2.**
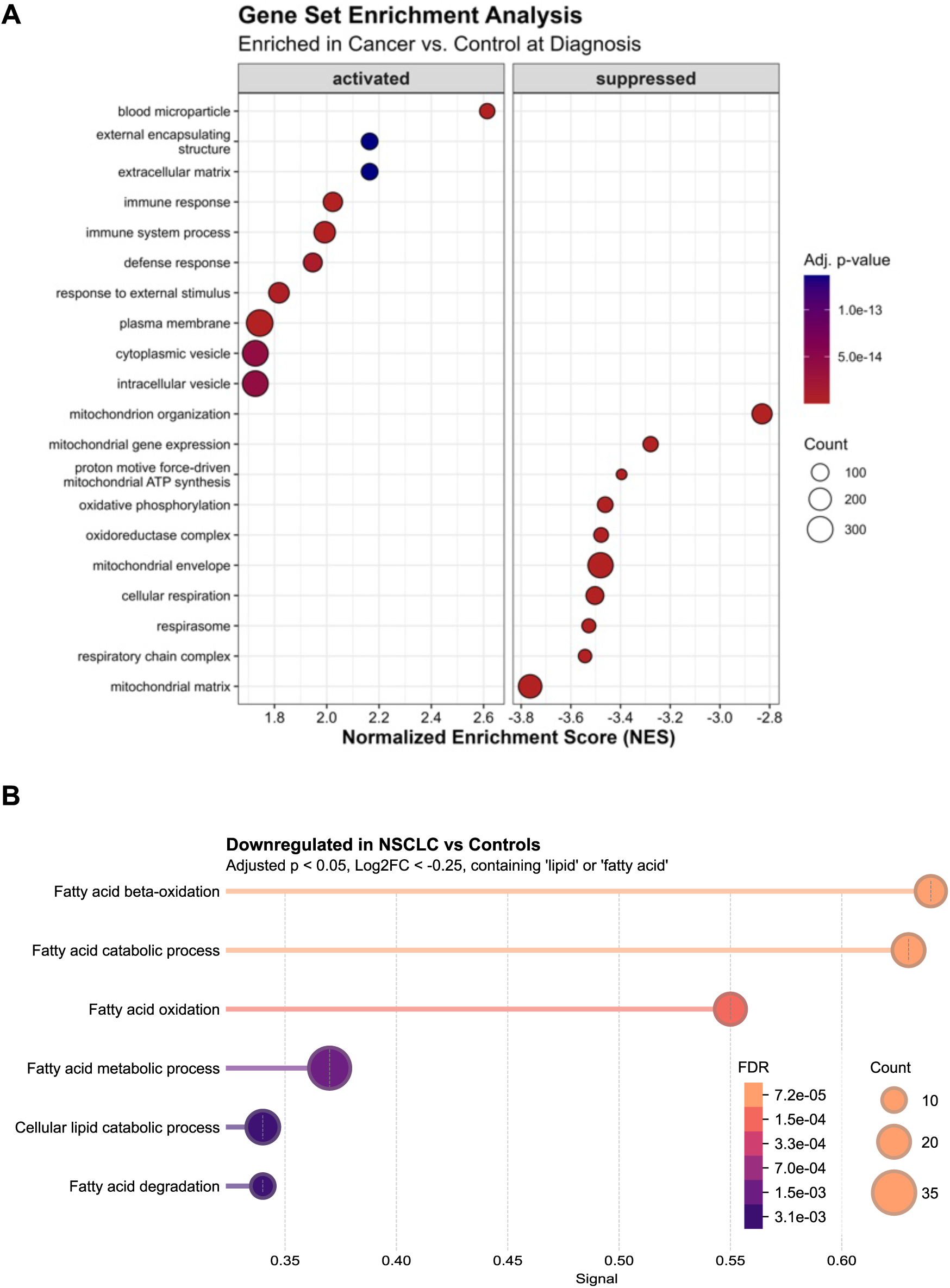
Additional proteomic enrichment analyses in skeletal muscle from patients with NSCLC. (A) Gene set enrichment analysis of proteins increased or decreased in NSCLC skeletal muscle compared with controls. (B) Over-representation analysis of proteins with adjusted P < 0.05 and log₂FC < −0.25 containing lipid- or fatty acid-related annotations. Enriched terms are shown as bubble plots, with bubble size indicating protein count and color indicating FDR.

**Supplementary Figure 3.**
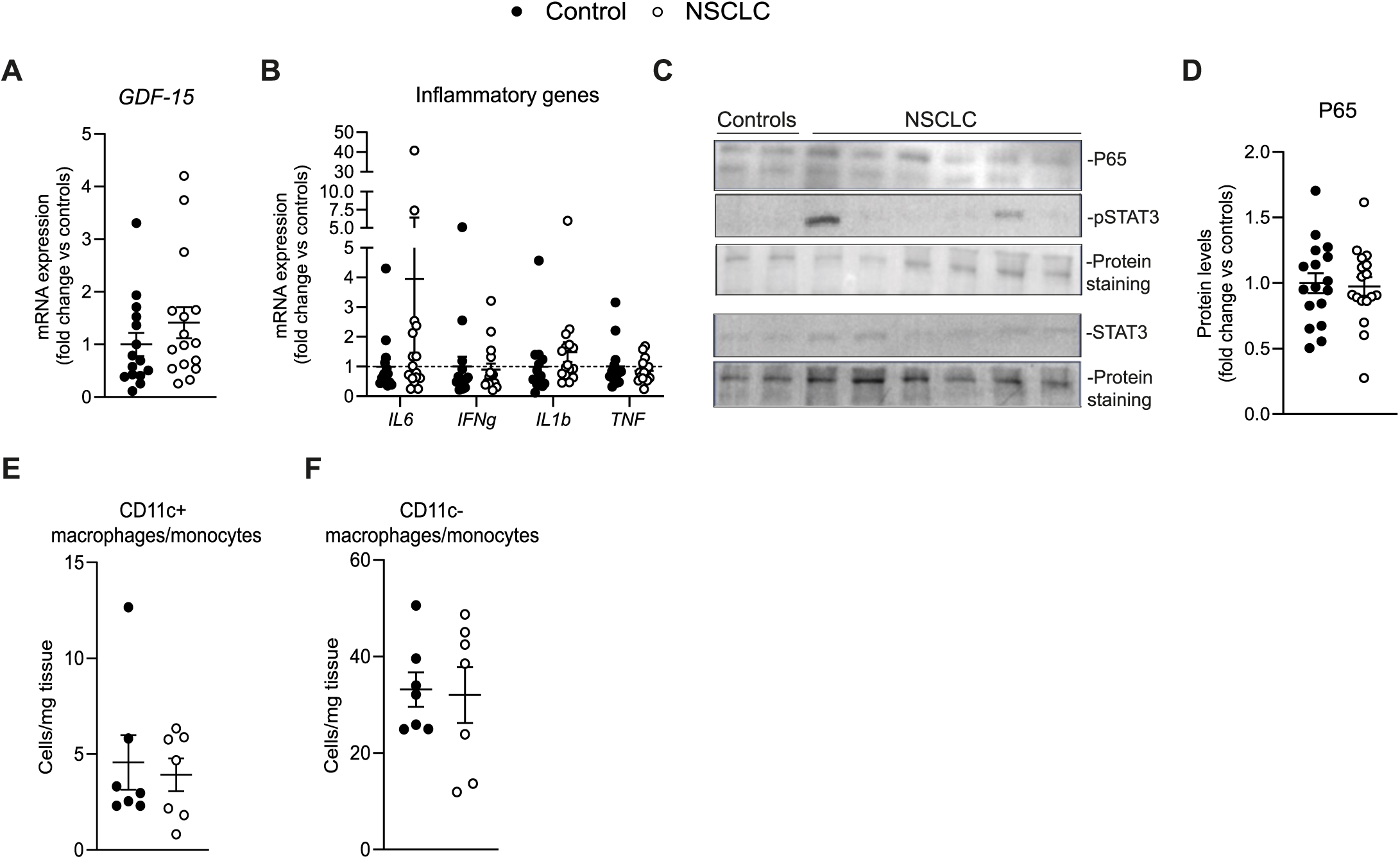
Supporting inflammatory signaling and immune-cell analyses in skeletal muscle from patients with NSCLC. (A) Skeletal muscle GDF-15 mRNA expression. (B) Skeletal muscle inflammatory gene expression. (C) Representative immunoblots of P65, pSTAT3, and STAT3, with total protein staining used as loading control. (D) Quantification of P65 protein abundance. (E,F) CD11c⁺ and CD11c⁻ monocyte/macrophage content (CD45⁺CD14⁺; cells per mg tissue). (A,D–F) Student’s t-test. (B) Two-way ANOVA.

**Supplementary Figure 4.**
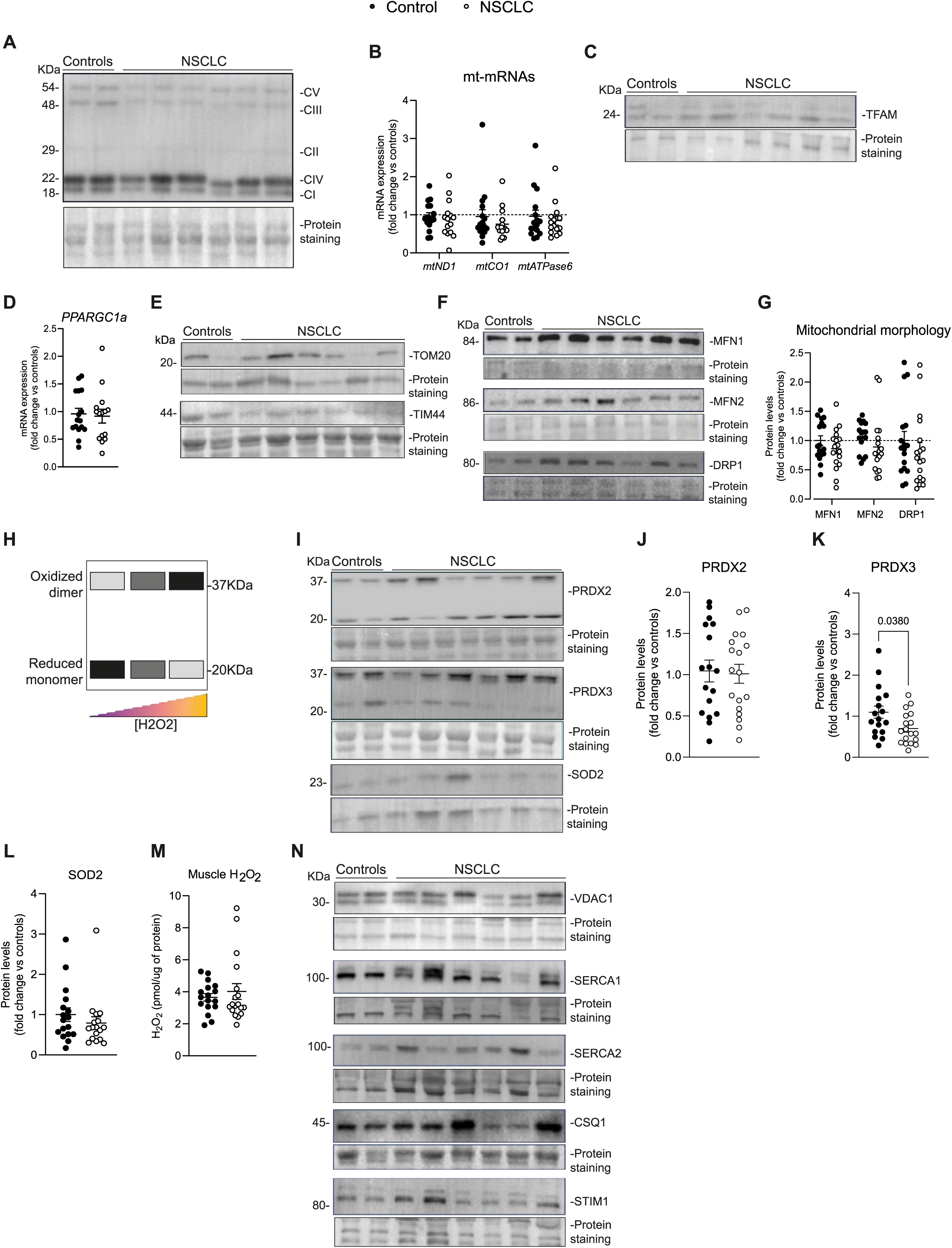
Supporting mitochondrial, redox, and calcium-handling analyses in skeletal muscle from patients with NSCLC. (A) Representative immunoblots of OXPHOS subunits, with total protein staining used as loading control. (B) Mitochondrial transcript abundance. (C) Representative immunoblot of TFAM. (D) PPARGC1a mRNA expression. (E) Representative immunoblots of TOM20 and TIM44. (F) Representative immunoblots of MFN1, MFN2, and DRP1. (G) Quantification of mitochondrial morphology-related proteins MFN1, MFN2, and DRP1. (H) Schematic representation of peroxiredoxin oxidation and dimerisation. (I) Representative immunoblots of PRDX2, PRDX3, and SOD2. Quantification of total PRDX2 (J), PRDX3 (K), and SOD2 (L) protein abundance. (M) Total skeletal muscle H₂O₂ levels. (N) Representative immunoblots of VDAC1, SERCA1, SERCA2, CSQ1, and STIM1, with total protein staining used as loading control. (B,G) Two-way ANOVA. (J–M) Student’s t- test.

**Supplementary Figure 5.**
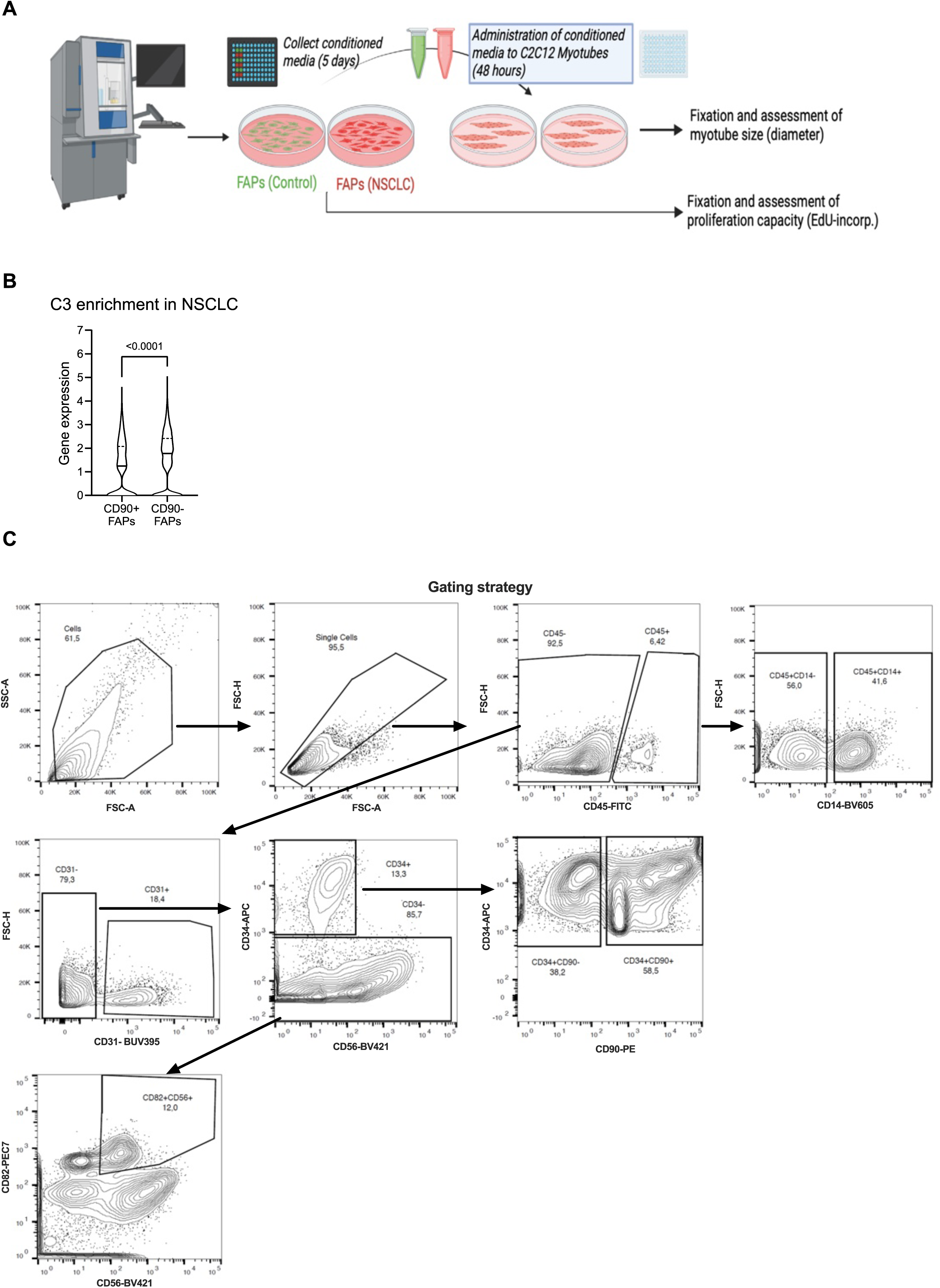
FAP conditioned-media workflow and C3 expression in FAP subpopulations. (A) Overview of in vitro experiments using FAPs isolated from controls and patients with NSCLC. Proliferative capacity was assessed by 5-ethynyl-2’-deoxyuridine (EdU) incorporation, and the effect of FAP-conditioned media on muscle size regulation was evaluated in C2C12 myotubes. (B) C3 expression in CD90⁺ and CD90⁻ FAP subpopulations. (C) Flow cytometry gating strategy used to identify FAP and MuSC populations.

## Supplementary Information

**Supplementary Table 1.** Primary antibodies used in immunoblotting assays.

| Targeted protein | Catalog no. | Company | Dilution |
| --- | --- | --- | --- |
| Anti-ATROGIN1 | sc-166806 | Santa Cruz Biotechnology | 1:1000 in TBST |
| Anti-MuRF1 | sc-398608 | Santa Cruz Biotechnology | 1:1000 in TBST |
| Anti-P62 (SQSTM1) | 51143 | Cell Signaling | 1:1000 in 2% skim milk |
| Anti-OXPHOS cocktail | ab110411 | Abcam | 1:1000 in 3% BSA |
| Anti-TFAM | ab131607 | Abcam | 1:1000 in 3% BSA |
| Anti-TOM20 | sc-17764 | Santa Cruz Biotechnology | 1:1000 in 2% skim milk |
| Anti-TIM44 | sc-390755 | Santa Cruz Biotechnology | 1:1000 in 2% skim milk |
| Anti-VDAC1 | sc-390996 | Santa Cruz Biotechnology | 1:1000 in 2% skim milk |
| Anti-MFN1 | sc-166644 | Santa Cruz Biotechnology | 1:1000 in TBST |
| Anti-MFN2 | 119255 | Cell Signalling | 1:1000 in TBST |
| Anti-DRP1 | sc-271583 | Santa Cruz Biotechnology | 1:1000 in TBST |
| Anti-PRDX2 | ab109387 | Abcam | 1:1000 in 3% BSA |
| Anti-PRDX3 | ab73349 | Abcam | 1:1000 in 3% BSA |
| Anti-SOD2 | o6984 | Milipore | 1:1000 in 3% BSA |
| Anti-P65 | sc-8008 | Santa Cruz Biotechnology | 1:1000 in 2% skim milk |
| Anti-STAT3 | Sc-8019 | Santa Cruz Biotechnology | 1:1000 in 3% BSA |
| Anti-p-STAT3(Tyr705) | 9138s | Cell Signalling | 1:1000 in 3% BSA |
| Anti-SERCA1 | MA3-912 | Invitrogen | 1:1000 in TBST |
| Anti-SERCA2 | MA3-910 | Invitrogen | 1:1000 in 2% skim milk |
| Anti-STIM1 | 49163 | Cell Signalling | 1:1000 in TBST |
| Anti-CSQ1 | PA1-193 | Invitrogen | 1:1000 in 2% skim milk |
| Anti-MHC (MF20) | MAB4470 | R&D | 1:25 in 1% BSA |

TBST: tris-buffered saline with tween 20; BSA: bovine serum albumin.

**Supplementary Table 2.**
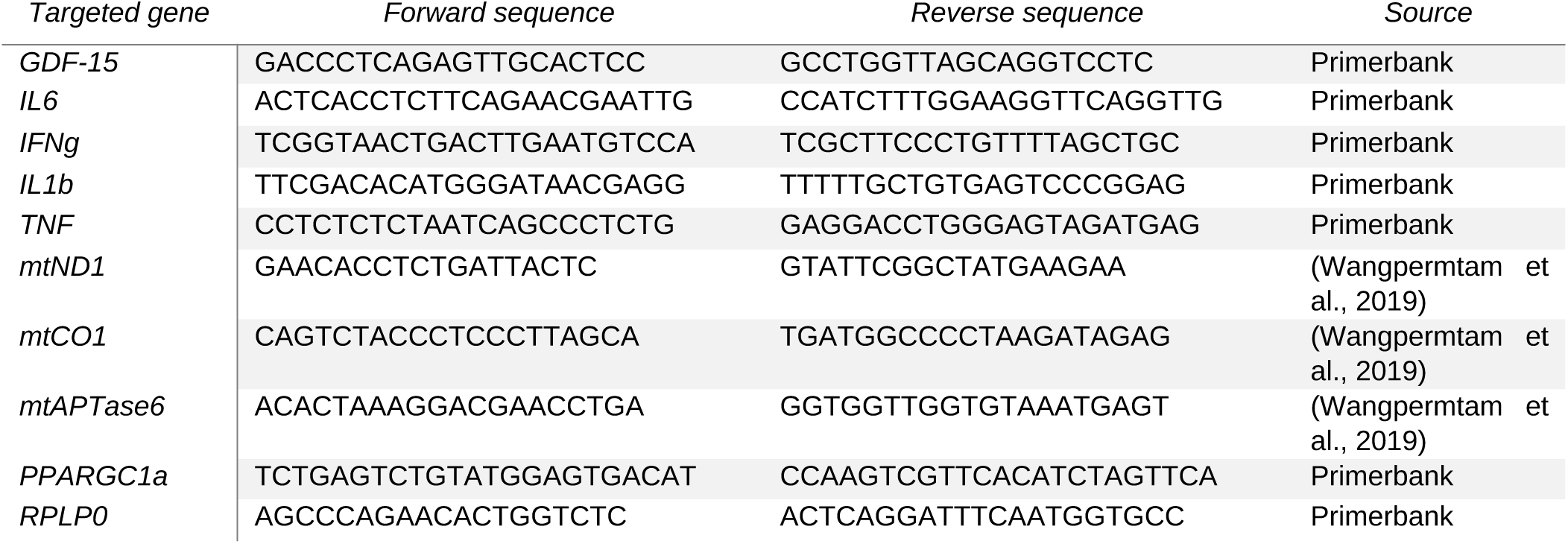
SYBR green primer pairs used in gene expression analyses.

**Supplementary Table 3.**
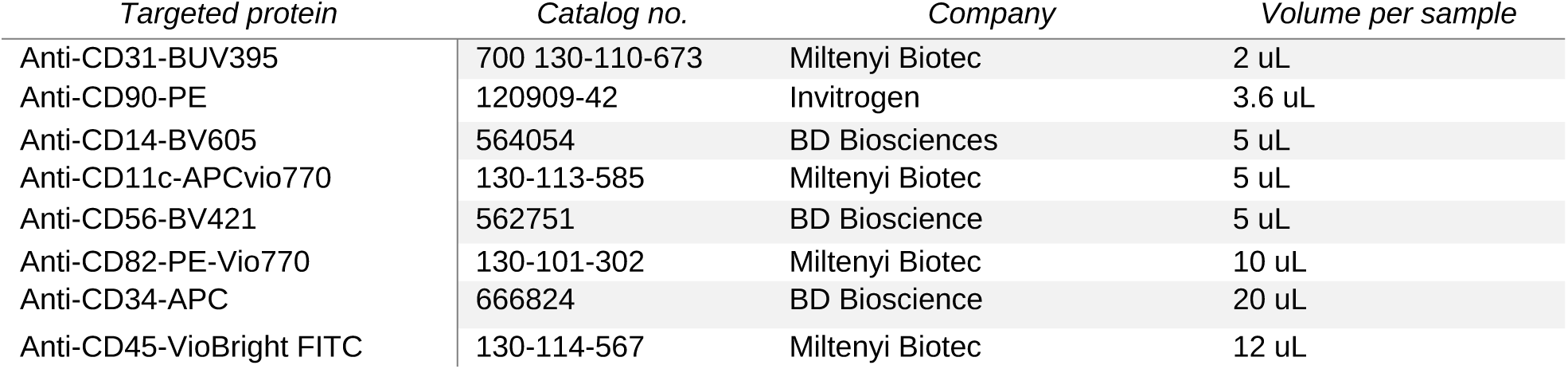
Primary antibodies used in fluorescence-activated cell sorting analyses.

## Notes

### Competing Interest Statement

The authors have declared no competing interest.

### Summary of Updates

We restructures the manuscript and included new data om FAPs

